# Task-induced neural covariability as a signature of approximate Bayesian learning and inference

**DOI:** 10.1101/081661

**Authors:** Richard D. Lange, Ralf M. Haefner

## Abstract

Perception can be characterized as an inference process in which beliefs are formed about the world given sensory observations. The sensory neurons implementing these computations, however, are classically characterized with firing rates, tuning curves, and correlated noise. To connect these two levels of description, we derive expressions for how inferences themselves vary across trials, and how this predicts task-dependent patterns of correlated variability in the responses of sensory neurons. Importantly, our results require minimal assumptions about the nature of the inferred variables or how their distributions are encoded in neural activity. We show that our predictions are in agreement with existing measurements across a range of tasks and brain areas. Our results reinterpret task-dependent sources of neural covariability as signatures of Bayesian inference and provide new insights into their cause and their function.

**Highlights:**

- General connection between neural covariability and approximate Bayesian inference based on variability in the encoded posterior density.
- Optimal learning of a discrimination task predicts top-down components of noise correlations and choice probabilities in agreement with existing data.
- Differential correlations are predicted to grow over the course of perceptual learning.
- Neural covariability can be used to ‘reverse-engineer’ the subject’s internal model.

## Introduction

Perceiving and acting in the world are remarkable feats of neural computation. A central goal of neuroscience is to simultaneously characterize both the neural mechanisms of these processes and, more abstractly, the computations implemented by those mechanisms (Marr, 1982). Currently, neural and computational levels of description lack clear links, even in such controlled settings as binary perceptual decision-making tasks (Parker and Newsome, 1998; Gold and Shadlen, 2007): neural models of perceptual decision-making are typified by encoding/decoding models built on population firing rates (Dayan and Abbott, 2001), while computational approaches typically model perception as approximate Bayesian inference (Knill and Pouget, 2004). This paper derives an analytical link between these frameworks, thus providing a novel explanation for observed changes in noise correlations due to factors such as task-switching and learning (Cohen and Newsome, 2008; Rabinowitz et al., 2015; Bondy et al., 2018; Ni et al., 2018).

The encoding/decoding framework models perceptual decision-making as a signal-processing problem: sensory neurons transform input signals, and downstream areas separate task-relevant signals from noise (Parker and Newsome, 1998). Theoretical arguments have shown how both encoded information (Zohary et al., 1994; Oram et al., 1998; Averbeck et al., 2006; Ecker et al., 2011; Moreno-Bote et al., 2014) and correlations between neurons and behavior (“choice probabilities”) (Shadlen et al., 1996; Haefner et al., 2013; Pitkow et al., 2015) depend on correlations among pairs of neurons, motivating numerous experimental studies into the nature of so-called “noise correlations” (Cohen and Newsome, 2008; Bondy et al., 2018; Goris et al., 2014; Ecker et al., 2014; 2016; Pitkow et al., 2015) (reviewed in (Kohn et al., 2016)). However, the extent to which choice probabilities and noise correlations are due to causally feedforward or feedback mechanisms is largely an open question (Nienborg and Cumming, 2009; Bondy et al., 2018; Goris et al., 2014; Wimmer et al., 2015) that has profound implications for their computational role (Nienborg and Cumming, 2010; Kohn et al., 2016; Lange and Haefner, 2017; Lueckmann et al., 2018; Macke and Nienborg, 2019).

The Bayesian inference framework, on the other hand, premises that the goal of sensory systems is to infer the latent causes of sensory signals (von Helmholtz, 1925) (Figure 1). This has motivated numerous theories of neural coding in which neural activity represents distributions of inferred variables (Zemel et al., 1998; Knill and Pouget, 2004; Fiser et al., 2010; Pouget et al., 2013; Ma and Jazayeri, 2014; Gershman and Beck, 2016). Bayesian inference further provides a rationale for the preponderance of feedback connections in the brain, which have been hypothesized to communicate contextual prior information or expectations (Mumford, 1992; Lee and Mumford, 2003; Summerfield and de Lange, 2014; de Lange et al., 2018).

**Figure 1.**
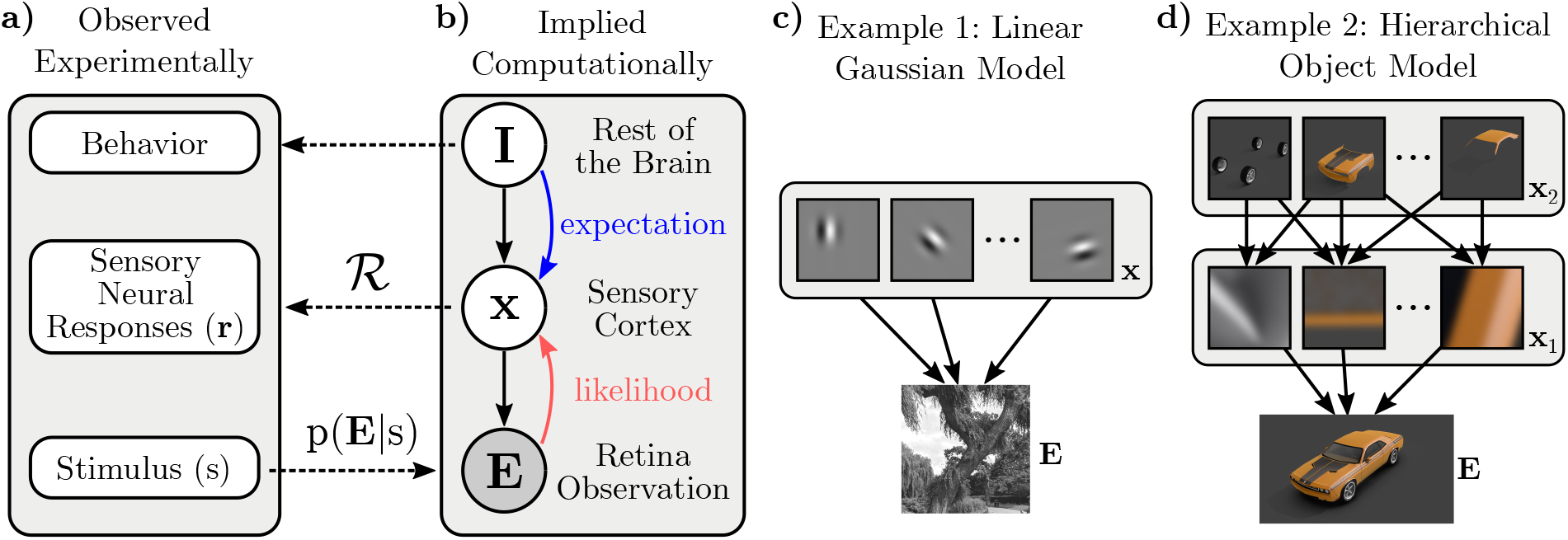
Illustration of the components of our framework and how they relate to experimentally observed quantities. **a-b)** The experimenter varies the sensory evidence, **E**, (e.g. images on the retina) according to s (e.g. orientation). The brain computes p(**x**,**I**|**E**), its beliefs about latent sensory variables of interest conditioned on those observations. **I** represents other “internal state” variables that are probabilistically related to **x**. The recorded neurons are assumed to encode the brain’s posterior beliefs about **x** through a distributional representation scheme, 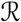. In the case of perceptual discrimination tasks, behavior is used to infer “categorical beliefs” about the stimulus, which are a subset of **I**. Solid black arrows represent statistical dependencies in the implicit generative model, not information flow. Dashed lines cross levels of abstraction. **c)** Example Generative Model 1: Olshausen and Field (1996) proposed that V1 performs inference in a linear-Gaussian “sparse coding” model fit to natural images. Here, **x** would correspond to the intensities of the Gabor elements in a given image. **d)** Example Generative Model 2: along the ventral stream, object recognition has been hypothesized to invert a generative model which proceeds from objects to parts to image features to images. **x** corresponds to inferred features at any level.

Here, we provide a missing link between these two frameworks: we show how principles of probabilistic learning and inference predict both task-dependent changes in the correlated variability among neural responses and the relationship between those responses and behavior. Assuming that neural responses represent posterior beliefs in a generative model of sensory inputs (von Helmholtz, 1925; Lee and Mumford, 2003; Kersten et al., 2004; Fiser et al., 2010), we derive predictions for how causally feedback or top-down components of neurons’ choice probabilities and noise correlations should depend on the neurons’ tuning to a stimulus.

Surprisingly, we find that after learning a task, the key signature of approximate inference in sensory responses are so-called “differential” or “information-limiting” correlations (Moreno-Bote et al., 2014). As a direct corollary, we predict these correlations to increase during task-learning. We further suggest a new way to interpret low-dimensional variability and choice probabilities in sensory neural populations as signatures of varying beliefs fed back to sensory areas. These results explain puzzling task-dependent patterns of noise correlations reported in previous studies (Cohen and Newsome, 2008; Rabinowitz et al., 2015; Bondy et al., 2018; Haimerl et al., 2019). Finally, these results imply, conversely, that sensory neural data can be used to infer a subject’s beliefs in a task, which we illustrate in simulation. Our results provide a normative justification for the growing empirical evidence for task-and choice-dependent feedback to sensory areas – which is hard to justify in the classic framework – by re-interpreting this feedback as a signature of a broad class of hierarchical inference algorithms.

## Results

Our results are organized as follows: first, we relate general distributional neural codes to neural tuning curves and correlated variability. We then apply this framework to the case of two-alternative forced-choice tasks and show that, after learning, trial-by-trial variations in a subject’s categorical beliefs imply noise correlations previously described as “differential” or “information-limiting”. We then generalize these results to incorporate task-independent noise. These results predict clear signatures of Bayesian inference and learning in pairwise neural firing rate statistics, which we compare with existing data. Finally, we illustrate how observed neural correlations can be used, conversely, to infer a subject’s internal beliefs from neural responses.

### Sources of neural variability in distributional codes

Following previous work, we assume that the brain has learned an implicit generative model of its sensory inputs (Figure 1c-d) (Lee and Mumford, 2003; Fiser et al., 2010; Olshausen and Field, 1997; Kersten et al., 2004; Yuille and Kersten, 2006), and that populations of sensory neurons encode posterior beliefs over latent variables in the model conditioned on sensory observations: a hypothesis we refer to as “posterior coding.” The responses of such neurons necessarily depend both on information from the sensory periphery, and on relevant information in the rest of the brain. In a hierarchical model, likelihoods are computed based on feedforward signals from the periphery, and contextual expectations are relayed by feedback from other areas (Lee and Mumford, 2003) (Figure 1b).

In our notation, **E** is the variable directly observed by the brain – the sensory input or evidence – and **x** is the (typically high-dimensional) variable whose posterior is assumed to be represented by the recorded neural population under consideration. **I** is a high-dimensional vector representing all other internal variables in the brain that are probabilistically related to, and hence determine “expectations” for **x** (Figure 1b)^1^. For instance, when considering the responses of a population of V1 neurons, **E** is the image on the retina, and **x** has been hypothesized to represent the presence or absence of Gabor-like features at particular retinotopic locations (Bornschein et al., 2013) or the intensity of such features (Olshausen and Field, 1996; Schwartz and Simoncelli, 2001) (Figure 1c), though our results are independent of the exact nature of **x**. In higher visual areas, variables could be related to the features or identity of objects and faces (Kersten et al., 2004; Yuille and Kersten, 2006) (Figure 1d). **I** represents higher-level variables, as well as knowledge about the visual surround, task-related knowledge about the probability of upcoming stimuli, etc.

The rules of Bayesian inference allow us to derive expressions for variability in posterior distributions as the result of learning and inference. Importantly, the rules of Bayesian inference apply to computational variables (Figure 1b); it is a conceptually distinct step to link variability in posteriors to variability in neurons encoding those posteriors. We use 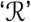 to denote the encoding from distributions over internal variables **x** into neural responses (Figure 2a,b). For reasonable encoding schemes 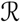, the chain rule from calculus applies: small changes in the encoded posterior result in small changes in the expected statistics of neural responses (Figure 2c, Methods). For instance, we can nexpress the change of a single neuron’s firing rate, *f*, in response to a change in stimulus, *s*, as

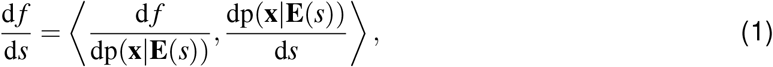

where 〈·,·〉 is an inner product in the space of distributions over **x**.^2^ The second term in brackets is the change in the posterior as s changes, and the first term relates those changes in the posterior to changes in the neuron’s firing rate.

**Figure 2.**
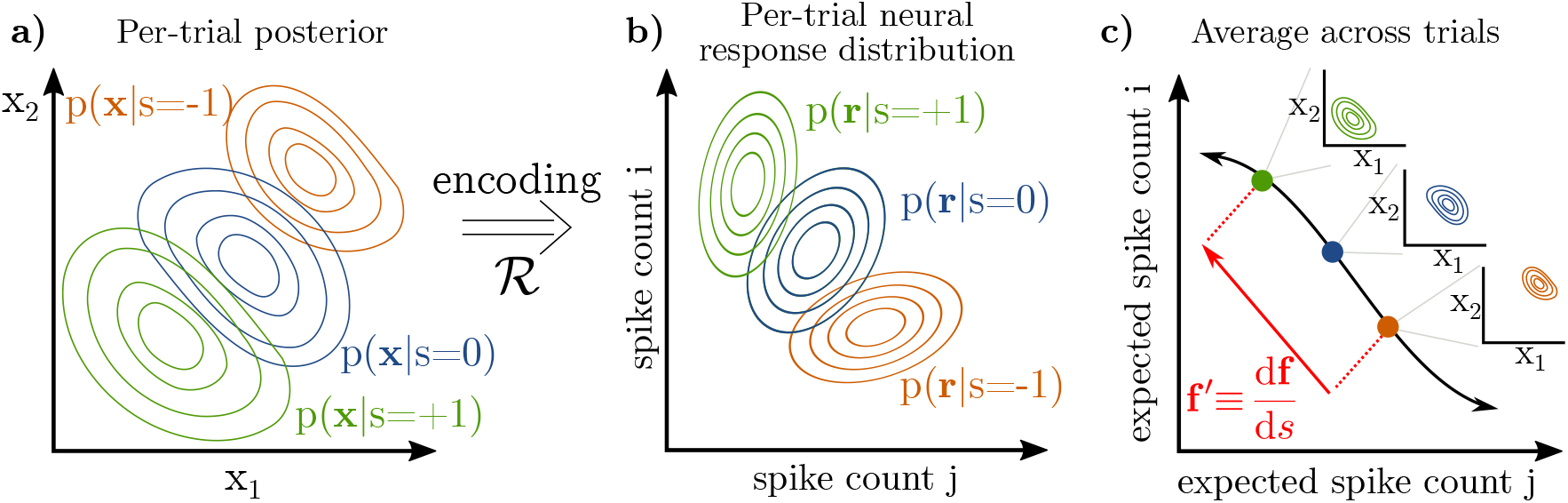
Neural representation of probability distributions. **a-b)** If neural responses encode a distribution over latent variables **x**, then one may think of the relation between **x** and **r** as a mapping from the space of distributions of latent variables (a) to the space of distributions of neural responses (b). Any given distribution on **x** may be stochastically encoded in **r**, for instance by Monte Carlo samples or by noisily representing parameters. Our derivation assumes that smoothly changing posteriors (a) corresponds to smooth changes in neural responses (b). **c)** Mean spike counts (or firing rates) across trials define a tuning curve. **f′** is the tangent vector to the tuning curve. It encodes, in part, the change in the underlying posterior over **x** (insets).

It follows that there are two sources of neural variability acting at different levels of abstraction: variability in the encoding of a given posterior (Figure 3a-c), and variability in the posterior itself (Figure 3d-f) (Beck et al., 2012).

**Figure 3.**
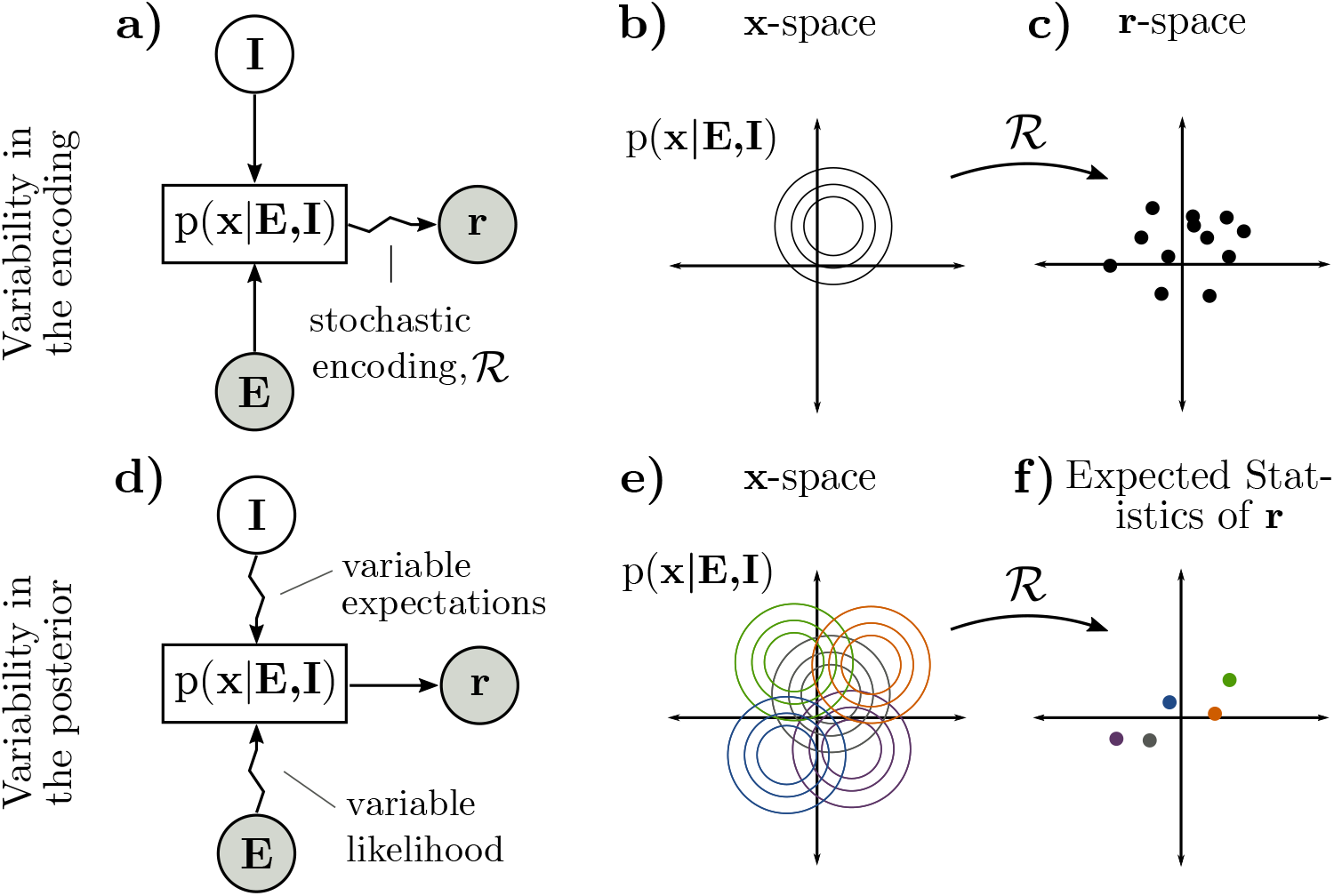
Neural co-variability may arise due to either (a-c) stochastic encoding or (d-f) variability in the posterior. **a)** Consider the case where there is no variability in **I** or **E** and inference is exact, but posteriors are noisily realized in neural responses **r**. **b)** Exact inference always produces the same posterior for **x** for fixed **E** and **I**. **c)** The neural encoding of a given distribution may be stochastic, so a single posterior (b) becomes a distribution over neural responses **r**. The shape of this distribution may or may not relate to the shape of the posterior in (b), depending on the encoding (e.g. there is a correspondence in sampling, but not in parametric codes). **d)** Noise perturbs the likelihood, and the subject’s beliefs vary. Both affect the posterior. Variable beliefs are the subject of our initial results, while noise will be considered later. **e)** Variability in the posterior can be thought of as a distribution over the space of possible posteriors. **f)** Each individual posterior in (e) is a point in the space of expected statistics of **r**, such as expected spike counts. Variability in the underlying posterior may appear as correlated variability in spike counts.

Distributional coding schemes (Zemel et al., 1998;Fiser et al., 2010;Pouget et al., 2013;Gershman and Beck, 2016) typically assume that a given posterior may be realized in a distribution of possible neural responses, which we refer to as **variability in the encoding** (Figure 3a-c). For instance, it has been hypothesized that neural activity encodes samples stochastically drawn from the posterior (Hoyer and Hyvärinen, 2003; Buesing et al., 2011; Pecevski et al., 2011; Savin and Denève, 2014; Petrovici et al., 2016; Haefner et al., 2016; Aitchson and Lengyel, 2016; Orbán et al., 2016; Aitchison et al., 2018). Alternatively, neural activity may noisily encode parameters of an approximate posterior (Ma et al., 2006; Beck et al., 2008; 2011; 2013; Raju and Pitkow, 2016; Pitkow and Angelaki, 2017; Vertes and Sahani, 2018). Such distributional encoding schemes are reviewed in (Fiser et al., 2010; Pouget et al., 2013; Gershman and Beck, 2016). Previous work has linked (co)variability in neural responses to sampling-based encoding of the posterior (Hoyer and Hyvärinen, 2003; Berkes et al., 2011; Orbán et al., 2016; Haefner et al., 2016; Bányai et al., 2019; Bányai and Orbán, 2019). Our results are complementary to these; here we study trial-by-trial changes in the posterior itself, and how these changes affect the expected statistics of neural responses such as mean spike count and noise correlations of neural responses. Importantly, our results apply to a wide class of distributional codes including all of the above (Methods).

To a first approximation, trial-by-trial **variability in the encoded posterior** manifests as neural (co)variability that simply sums with the variability in the encoding already discussed (Figure 3d-f). For instance, noise in the stimulus, sensory measurements, and afferent neural signals affect the likelihood (Faisal et al., 2008; Stocker and Simoncelli, 2006; Körding et al., 2007), and variable internal states may influence sensory expectations through feedback (Nienborg and Roelfsema, 2015; Lange and Haefner, 2017). We will ignore such task-independent noise for our initial results. Instead, our first results concern variability in the posterior due to variability in task-relevant beliefs or expectations (Nienborg and Roelfsema, 2015; Haefner et al., 2016). Variable expectations may reflect a stochastic approximate inference algorithm (Hoyer and Hyvärinen, 2003) or model mismatch, for example if the brain picks up on spurious dependencies in the environment as part of its model (Beck et al., 2012; Yu and Cohen, 2009; Fründ et al., 2014; Fischer and Whitney, 2014). In the remainder of this paper, we make these ideas explicit for the case of two-alternative decision-making tasks for which much empirical data exists.

### Inference and discrimination with arbitrary sensory variables

In the special case of inference in a two-alternative discrimination task, stimuli are parameterized along a single dimension, s, and subjects learn to make categorical judgments according to an experimenter-defined boundary which we assume is at s = 0 (Figure 4a). We will use *C* ∈ {1, 2} to denote the two categories, corresponding to *s* < 0 and *s* > 0. Throughout this paper, our running example will be of orientation discrimination, in which case *s* is the orientation of a grating with *s* = 0 corresponding to horizontal, and C refers to clockwise or counter-clockwise tilts (Figure 4b). While our derivations make no explicit assumptions about the nature of the brain’s latent variables, **x**, our illustrations will use the example of oriented Gabor-like features in a generative model of images (Figure 1c, Figure 4b).

**Figure 4.**
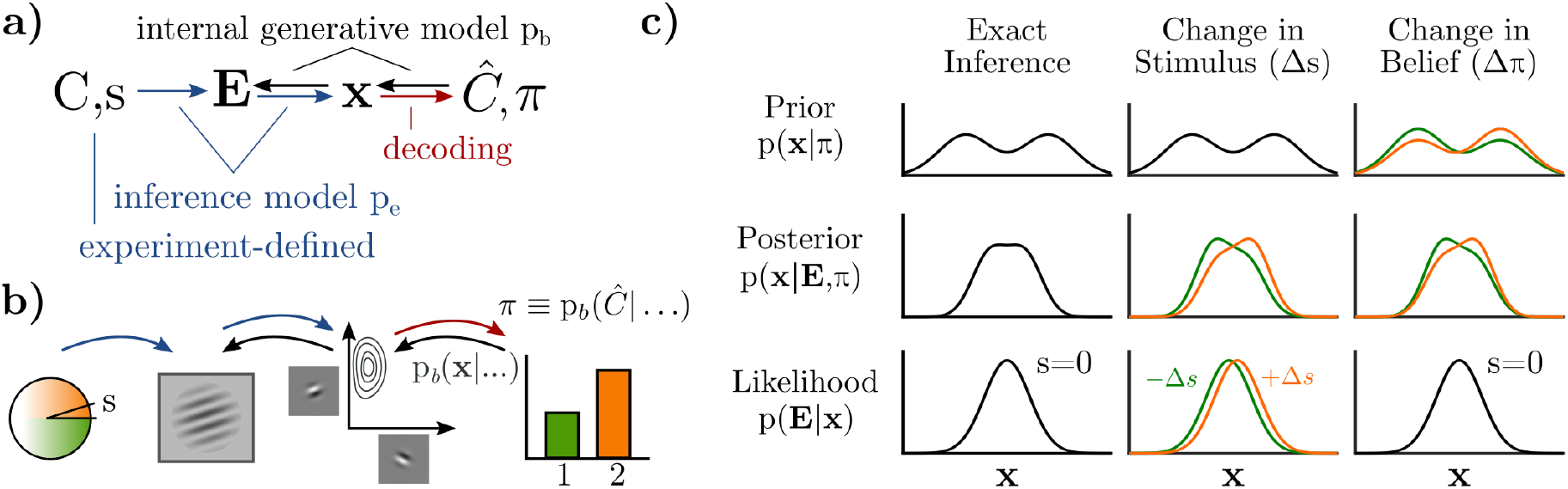
**a)** A discrimination task defines a joint distribution between category *C* and stimulus parameter s, which gives rise to sensory inputs **E**. The brain performs inference over sensory latent variables (**x**) and estimated category (*Ĉ*) conditioned on the stimulus (**E**). Graded beliefs about the binary category are expressed as *π* ≡ p_b_(*Ĉ*|…). Implicitly, these inferences are with respect to an internal model pb (black arrows). A Bayesian observer learns a joint distribution between **x** and *Ĉ*, implying bi-directional influences during inference: **x** → *Ĉ* is analogous to “decoding,” while *Ĉ* → **x** conveys task-relevant expectations. **b)** Conceptual illustration of (a) for fine orientation discrimination, where latents **x** are Gabor-like features in a generative image model. The “decoder” then forms a belief, *π*, over internal estimates of the category. **c)** Visualization of how the prior (top row) and likelihood (bottom row) contribute to the posterior (middle row), with **x** as a one-dimensional variable. Changes to *s* change the likelihood (middle column). Changes in expectation, *π*, are changes in the prior (right column). Crucially, changes in the posterior in both cases (middle row) are approximately equal.

Whereas much previous work on perceptual inference assumes that the brain explicitly infers relevant quantities defined by the experiment (Gold and Shadlen, 2007; Knill and Pouget, 2004; Ma et al., 2006; Beck et al., 2008), we emphasize the distinction between the external stimulus quantity being categorized, *s*, and the latent variables in the subject’s sensory model of the world, **x**. For the example of orientation discrimination, a grating image **E**(*s*) is rendered to the screen with orientation *s*, from which V1 infers an explanation of the image as a combination of Gabor-like basis elements, **x**. The task of downstream areas of the brain – which have no direct access to **E** nor *s* – is to estimate the stimulus category based on a probabilistic representation of **x** (Figure 4b) (Haefner et al., 2016; Shivkumar et al., 2018). Crucially it is the posterior over **x**, rather than over s, which we hypothesize is represented by sensory neurons.

### Task-specific expectations

Probabilistic relations are inherently bi-directional: any variable that is predictive of an other variable will, in turn, be at least partially predicted by that other variable. In the context of perceptual decision-making, this means that sensory variables, **x**, that inform the subjects’ internal belief about the category, *Ĉ*, will be reciprocally influenced by the subject’s belief about the category (Figure 4a). Inference thus gives a normative account for feedback from “belief states” to sensory areas: changing beliefs about the trial category entail changing expectations about the sensory variables whenever those sensory variables are part of the process of forming those beliefs (Lee and Mumford, 2003; Lee et al., 2014; Nienborg and Roelfsema, 2015; Haefner et al., 2016).

A well-known identity for well-calibrated probabilistic models is that their prior is equal to their average inferred posterior (Dayan and Abbott, 2001; Fiser et al., 2010; Berkes et al., 2011). We derive an analogous expression for the optimal prior over **x** upon learning the statistics of a task (Methods):

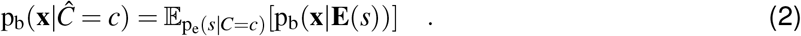

Equation (2) states that, given knowledge of an upcoming stimulus’ category, *Ĉ* = *c*, the optimal prior on **x** is the average posterior from earlier trials in the same category (Stocker and Simoncelli, 2007). The subscript ‘b’ refers the brain’s internal model, while ‘e’refers to the experimenter-defined model (Figure 4a, Methods). To use the orientation discrimination example, knowing that the stimulus is “clockwise” increases the expectation that more clockwise-tilted Gabor features will be present, since they were inferred to be present in earlier clockwise trials. Importantly, equation (2) is true regardless of the nature of **x** or *s*. It is a *self-consistency* rule between prior expectations and posterior inferences that is true for any ideal learner given sufficient experience (Dayan and Abbott, 2001; Berkes et al., 2011) (see also Supplemental Text). This self-consistency rule allows us to relate neural responses to the stimulus (*s*) to neural responses to internal beliefs (*π*) without specific assumptions about **x**.

In binary discrimination tasks, the subject’s belief about the correct category is a scalar quantity, which we denote by *π* = *p*(*Ĉ* = 1). Given *π*, the optimal expectations for **x** are a correspondingly graded mixture of the per-category priors:

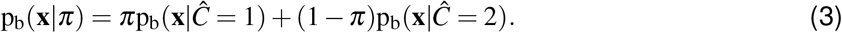

The posterior over **x** for a single trial depends on both the stimulus and belief for that trial:

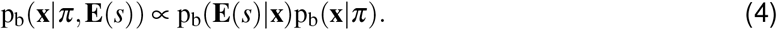

We will next derive the specific pattern of neural correlated variability when *π* varies.

### Variability in the posterior due to changing expectations

Even when the stimulus is fixed, subjects’ beliefs and decisions are known to vary (Parker and Newsome, 1998). Small changes in a Bayesian observer’s categorical belief (Δ*π*) result in small changes in their posterior distribution over **x**, which can be expressed as the derivative of the posterior with respect to *π* (assuming the stimulus has been fixed to the category boundary):

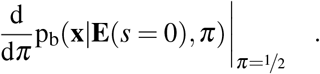

Our first result is that this derivative is *approximately proportional* to the derivative of the posterior with respect to the stimulus. Mathematically, the result is as follows:

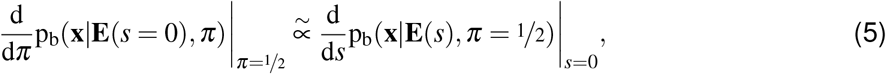

where the symbol 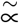 should be read as “approximately proportional to” (see Methods for proof) (Figures 4c, S2).

Equation (5) states that, for a Bayesian observer, small variations in the stimulus around the category boundary have the same effect on the inferred posterior over **x** as small variations in their categorical beliefs. The proof makes four assumptions: first, the subject must have fully learned the task statistics, as specified by equations (2) and (3). Second, the two stimulus categories must be close together, i.e. the task must be near or below psychometric thresholds, such that neural dependencies on the stimulus are approximately linear. Third, variations of stimuli within each category must be small. We further discuss these conditions and possible relaxations in the Supplemental Text. Finally, we have assumed that there are no additional noise sources causing the posterior to vary; we consider the case of noise in the section “Effects of task-independent noise” below.

### Feedback of variable beliefs implies differential correlations

Applying the “chain rule” in equation (1) to equation (5), it directly follows that

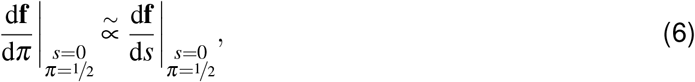

implying that the effect of small changes in the subject’s categorical beliefs (*π*) is approximately proportional to the effect of small changes in the stimulus on the responses of sensory neurons that encode the posterior. Both induce changes to the mean rate in the **f′** ≡ d**f**/d*s* direction. Because **f′** itself is task-dependent, variable task-relevant beliefs will add to neural covariability in the **f′** direction above and beyond whatever intrinsic covariability was present before learning. We obtain, to a first approximation, the following expression for the noise covariance between neurons *i* and *j*:

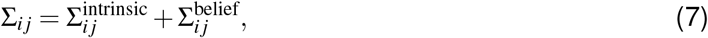

where 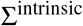 captures “intrinsic” noise such as Poisson noise in the encoding. It follows from (6) that

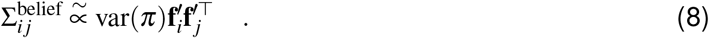

Interestingly, this is exactly the form of so-called “information-limiting” or “differential” covariability (Moreno-Bote et al., 2014). Whereas in the feedforward framework this covariability arises due to variability in the sensory inputs limiting the information about *s* in the population (Moreno-Bote et al., 2014; Kanitscheider et al., 2015; Kohn et al., 2016), here it arises due to feedback of variable beliefs about the stimulus category. Unless these beliefs are true, or unless downstream areas have access to and can compensate for *π*, the differential covariability induced by *π* limits information like its bottom-up counterpart (Kohn et al. (2016); Lange and Haefner (2017); Bondy et al. (2018); also see Discussion). Importantly, unlike feedforward differential covariability, the feedback differential covariability predicted here arises as the result of task-learning, which makes their relative strength an empirically decidable question.

### Variable beliefs imply structure in choice probabilities

A direct prediction of the feedback of beliefs *π* to sensory areas is that the average neural response preceding choice 2 will be biased in the +**f′** direction, and the average neural response preceding choice 1 will be biased in the −**f′** direction, since the subject’s actual choices will be based on their belief, *π*. Feedback of *π* will therefore introduce additional correlations between neural responses and choice above and beyond those predicted by a purely feedforward “readout” of the sensory neural responses (Parker and Newsome, 1998; Nienborg and Cumming, 2009; Nienborg et al., 2012; Haefner et al., 2013; Pitkow et al., 2015; Wimmer et al., 2015; Haefner et al., 2016). This top-down component of choice probability is predicted to be proportional to neural sensitivity:

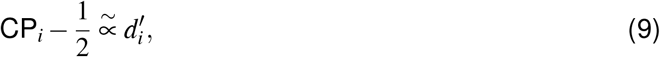

where 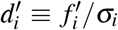 is the “d-prime” sensitivity measure of neuron *i* from signal detection theory (Green and Swets, 1966) (Figure 6a; Methods). Interestingly, the classic feedforward framework makes the same prediction for the relation between neural sensitivity and choice probability assuming an optimal linear decoder (Haefner et al., 2013; Pitkow et al., 2015), raising the question to what degree the empirically observed relationship between CPs and neural sensitivity (Law and Gold, 2008) is due to changes in the feedforward read-out over learning as commonly assumed (Parker and Newsome, 1998; Law and Gold, 2009) versus changes in feedback signals due to variable beliefs.

### Effects of task-independent noise

The above results assumed no measurement noise nor variability in other internal states besides the relevant belief *π*. In the presence of noise, the posterior itself changes from trial to trial even for a fixed stimulus *s* and fixed beliefs *π* (Stocker and Simoncelli, 2006). To study the consequences of this added variability, we introduce avariable, *ε*, that encompasses all sources of task-independent noise each trial, and condition the posterior on its value: p(**x**|**E**(*s*),*π*;*ε*) (Methods). This impacts our main results in two principal ways, laid out in the following two sections: first, although ideal learning still implies that the average posterior equals the prior (equation (2)), the “average” must now be taken over both *s* and the distribution of noise p(*ε*). Second, task-independent noise will interacts a task-dependent prior (Figure 5) which also has a task-dependent effect on neural covariability.

**Figure 5.**
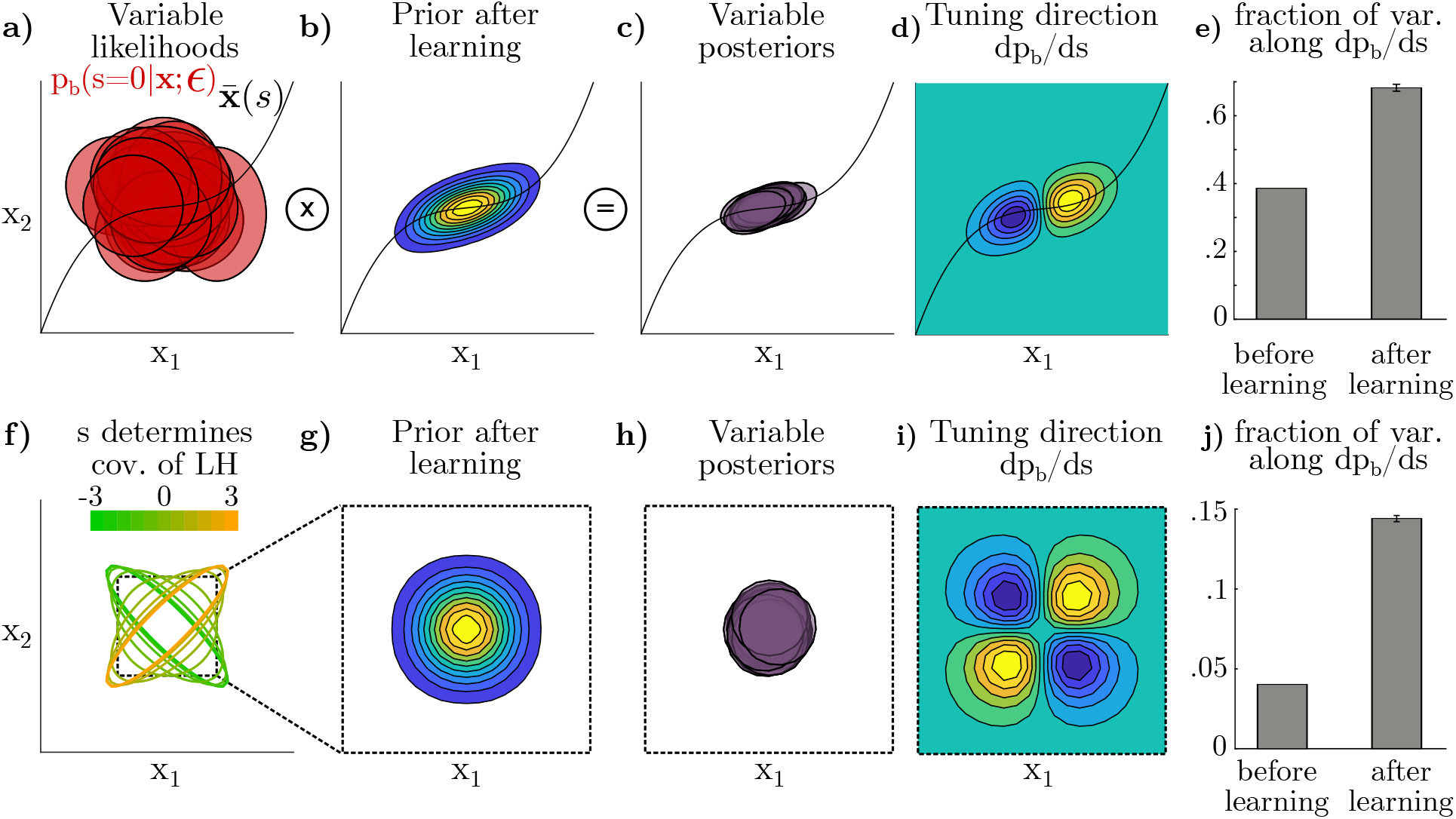
Sketch of how variable likelihoods both determine and interact with the shape of the prior. **a)** Visualization of task-independent variability producing a range of likelihoods with *s* = 0 fixed. For the first simulation, *s* parameterizes the mean of the likelihood along the curve 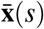. **b)** After learning, the prior is extended along 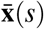, since it is the average of posteriors over all *s*. **c)** Posteriors in the zero-signal case, given by the product of the likelihoods in (a) with the prior in (b). **d)** The direction in this space corresponding to differential covariance in neurons is the dp_b_/d*s*-direction, averaged over instances of noise. **e)** The fraction of variance in posteriors (c) along the dp_b_/d*s*-direction. After learning, an larger fraction of the total variance is in the dp_b_/d*s*-direction. Error bars indicate ±1 standard deviation across runs. **f)** Whereas in (a)–(e) the external changes in *s* drove the *mean* of the likelihood, here we simulate changes to higher-order moments by keeping the mean of **x** fixed but parameterizing its shape with *s*, which has a uniform distribution in [−3,+3] (a.u.). Dashed inset indicates zoomed in plots in (g)–(i). **g-j)** as in (b)–(e) but using the likelihoods in (f). Dashed borders indicate zoooming to the box outlined in (f). While the overall magnitude of variance is smaller, the trend in (j) is analogous to (e): learning increases the fraction of variance in the dp_b_/d*s*-direction.

### Variable beliefs in the presence of noise

In the presence of noise, a neuron’s sensitivity to the stimulus, 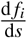, can be written as the *average* sensitivity of *f_i_* to changes in the posterior given *s*. On the other hand, a neuron’s sensitivity to feedback of beliefs, 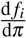, depends on the sensitivity of *f_i_* to the *average posterior* (Methods). Because the expected value of a function is not equal to the function of an expected value, the neural response to a change in belief (related to the average posterior) might therefore be different from the average neural response to a change in the stimulus, in general. However, there is a subclass of encoding schemes, 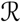, in which firing rates are linear with respect to *mixtures* of distributions over **x**. For those schemes the two expectations are therefore identical and we recover our earlier results for both task-dependent noise covariance (equation (8)) and structured choice probabilities (equation (9)) (Methods). We call these *Linear Distributional Codes* (LDCs). Examples of LDCs in the literature are given in the Discussion. We expect our results to degrade gracefully for codes that are nearly linear, or if the magnitude of the task-independent noise is small.

### Interactions between task-independent noise and task-dependent priors

Although we assumed that noise *ε* arises from task-independent mechanisms, it is nonetheless shaped by task learning:task-independent noise in the likelihood interacts with a task-specific prior to shape variability in the posterior (Figure 5). This implies a source of task-dependent correlation in neural responses representing a posterior that will be present even if a subject’s beliefs (*π*) do not vary. This idea is reminiscent of circuit models of the influence of task context on recurrent dynamics, shaping the manifold along which neural activity may feasibly vary (Huang et al., 2019; Doiron et al., 2016).

We again study the trial-by-trial variability in the posterior itself as opposed to the shape or moments of the posterior on any given trial. This can be formalized the covariance due to noise (ε) in the posterior *density* at all pairs of points **x**_*i*_, **x**_*j*_, i.e. Σ ≡ cov(p_b_(**x**_1_|…), p_b_(**x**_2_|…)). We show (Methods) that, to a first approximation, the posterior covariance is given by a product of the covariance of the task-independent noise in the likelihood, Σ^LH^(**x**_*i*_,**x**_*j*_), and the brain’s prior over **x**_*i*_ and **x**_*j*_:

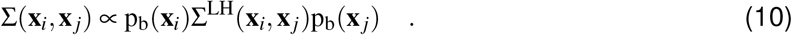

The effect of learning a task-dependent prior in equation (10) can be understood as “filtering” the noise, suppressing or promoting certain directions of variability in the space of posterior distributions. Differential correlations emerge from this process if variability in the dp_b_(**x**|…)/d***s***-direction is less suppressed than in other directions. Whether this is the case, and to what extent, depends on the interaction of *s* and **x**, an analytic treatment of which we leave for future work. Here, we present the results from two representative simulations, one in which the mean of **x** depends on *s* and one in which the covariance of **x** depends on *s*.

In both simulations, we assume **x** to be two-dimensional with isotropic Gaussian likelihoods over *s*. The prior was learned by iteratively applying equation (3), including noise, until convergence. Noise was added by jittering the mean and covariance of each likelihood (Figure 5a). In the first simulation, the *mean* of the likelihood non-linearly depended on *s* (Figure 5a-d). Small variations in *s* around the boundary *s* = 0 primarily translated the posterior, resulting in a two-lobed dp_b_/d*s* structure (Figure 5d). After learning, the prior sculpted the noise such that trial-by-trial variance in posterior densities was dominated by translations in the dp_b_(**x**|…)/d*s*-direction (Figure 5c+e).

The intuition behind this first simulation is as follows. During learning, both uninformative *s* = 0 and informative *s* < 0 or *s* > 0 stimuli are shown. As a result, the learned prior (equalling the average posterior) becomes elongated along the curve that defines the mean of the likelihood (Figure 5b), which is also the direction that defines dp_b_/d*s*. After learning, if noise shifts the likelihood along this curve, then the resulting posterior will remain close to that likelihood because the prior remains relatively flat along that direction. In contrast, noise that changes the likelihood in an orthogonal direction will be “pulled” back towards the prior. Thus, multiplication with the prior preferentially suppresses noise orthogonal to dp_b_/d*s*. Applying the chain rule from equation (1), this directly translates to privileged variance in the differential or **f′f′**^T^ direction in neural space.

To investigate whether this result only holds when the mean of the likelihood depends on the stimulus, we next held the mean of the likelihood constant and assumed that the stimulus is encoded in its (co)variance (Figure 5f). Otherwise, likelihoods, the learning procedure, and noise were identical to the first simulation. Interestingly, we again found that the variance in the dp_b_/d*s*-direction was enhanced relative to other directions after learning (Figure 5i-j), again implying differential correlations in the neural responses.

Note that whereas our results on variability due to the feedback of variable beliefs implied an increase in neural *covariance* along the **f′f′**^/T^-direction over learning, the effect of “filtering” the noise induces task-related noise *correlations* but does not necessarily increase nor decrease variance (depending on the brain’s prior at the initial stage of learning).

### Empirical hypothesis tests

To summarize, we have identified three signatures of Bayesian learning and inference: structured choice probabilities (equation (9)) and noise correlations (equation (8)) due to trial-by-trial feedback of beliefs *π*, and additional structure in noise correlations due to the “filtering” of task-independent noise. We emphasize that our results only describe how learning a task-specific prior *changes* these quantities, and makes no predictions about their structure before learning. Below we present five strategies to experimentally test our predictions and discuss their relation to existing empirical data.

First, our results predict that the top-down component of choice probability should be proportional to the vector of neural sensitivities to the stimulus (Figure 6a). Indeed, such a relationship between CP and *d* was found by many studies (reviewed in Nienborg et al. (2012)). However, this is only a weak test since this finding can also be explained in a purely feedforward framework (Law and Gold, 2009; Haefner et al., 2013), so the remaining strategies focus on predictions for correlated variability, which cannot be accounted for with feedforward mechanisms.

**Figure 6.**
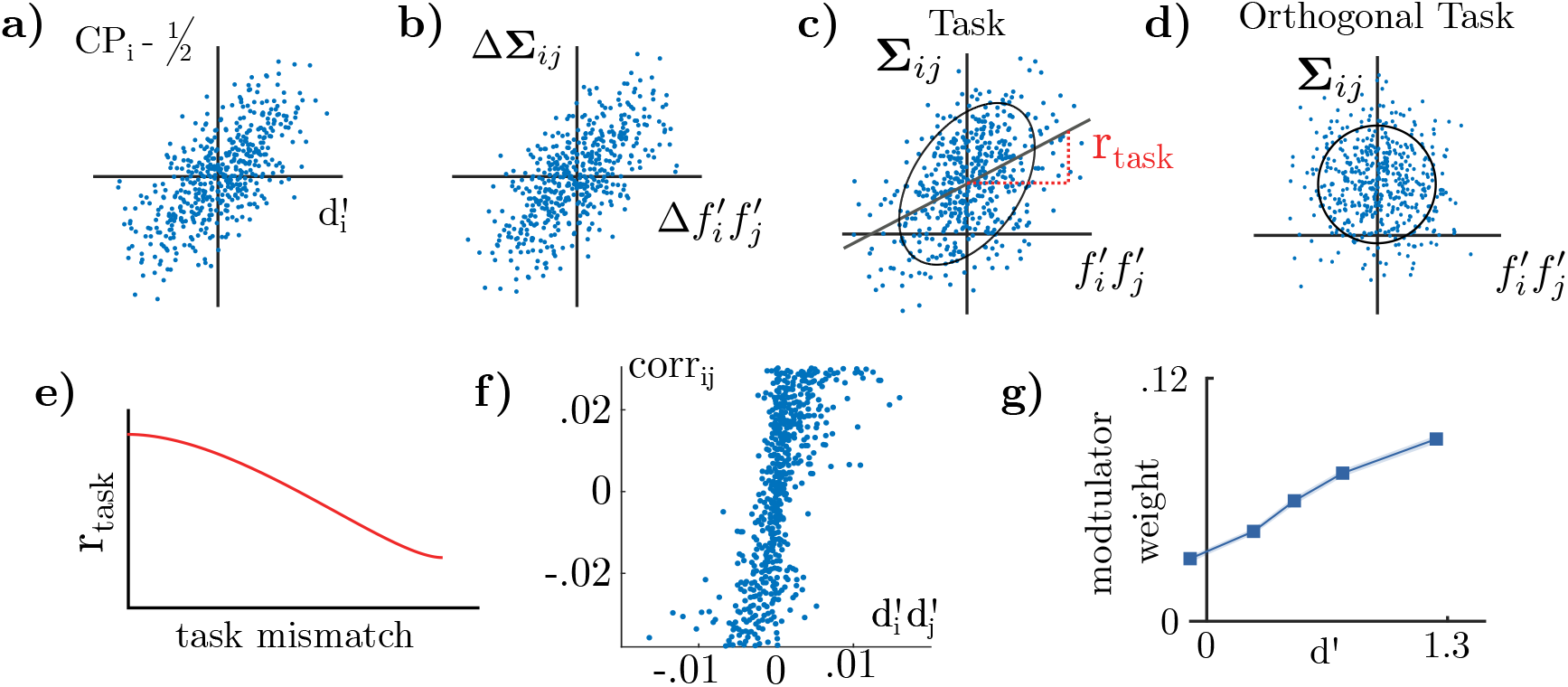
Predictions of the probabilistic inference framework. Σ denotes covariation, and corr denotes correlation. 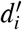 is the normalized sensitivity of neuron *i* defined as 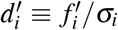. **a)** First prediction, in agreement with classical feedforward encoding-decoding models with optimal linear readout: neurons’ choice. probabilities should be proportional to their normalized sensitivity to the stimulus. **b)** Second prediction, requiring top-down signals: the difference in covariance structure between comparable tasks should be proportional to the difference in the product of tuning curve derivatives for each task. By subtracting out intrinsic covariability, this is a less noise-prone prediction than (c-e). **c)** Noise covariance induced by task-learning should be proportional to **f′f′**^T^. **d)** As a control, the relationship in (c) should not hold for neural sensitivities *d′* measured with respect to other tasks’ **f′** vectors. **e)** Summary of (c) and (d): *r_task_* should fall off when computed with respect to other hypothetical task directions (e.g. by predicting the **f′** vector for other tasks from tuning curves). f) Results of Rabinowitz et al. (2015) replotted, where it was found that the strength of top-down ‘modulator’ connections is linearly related to *d′*. **g)** Bondy et al. (2018) isolated the top-down, task-dependent component of noise correlations in macaque V1, and found a strong relation between elements of this correlation matrix and neural sensitivities (r = 0.61, p < 0.001, from original paper); similar to panel (b) divided by the standard deviation of neural responses.

A second strategy involves holding the stimulus constant while switching between two comparable tasks that a subject is performing, altering their task-specific expectations. The difference in neural response statistics to a stimulus that is *shared by both tasks* will isolate the task-dependent component to which the our predictions apply (Figure 6b). In this vein, Bondy et al. (2018) recorded from neural populations in macaque V1 while the monkeys switched between different coarse orientation tasks. They found that the changes in noise correlations were well-aligned with **d′d′**^T^ structure as predicted by equation (8) (Figure 6g). Note that a proportionality between covariance and **f′f′** is equivalent to a proportionality between correlation and **d′d′**. Cohen and Newsome (2008) recorded from pairs of neurons in area MT of two monkeys and found that correlations also changed as if caused by variability in internal belief (see Box 2 in Lange and Haefner (2017)). A critical requirement for this approach is that the stimulus distribution at *s* = 0 is matched between the two different tasks so that “intrinsic” covariability can be subtracted out (Methods).

A third, related, approach is to compare the amount of correlated variability in the current task’s direction with other “hypothetical” tasks as controls (Figure 6c-e). For instance in a coarse orientation discrimination task the covariability in the population response in the **f′**–direction of the actually performed task (e.g. vertical vs horizontal) should be larger than the variability in directions corresponding to other tasks (e.g. –45deg vs +45deg).

A fourth strategy is to *statistically* isolate the top-down component of neural variability within a single task using a sufficiently powerful regression model. Rabinowitz et al. (2015) used this type of approach to infer the primary top-down modulators of V4 responses in a change-detection task. They found that the two most important short-term modulators were closely aligned with the **f′**–direction corresponding to the monkey’s task (data replotted in Figure 6f).

Finally, our predictions can be tested through experimental manipulation of feedback pathways. In particular, we predict that the task-dependent **f′f′**^T^ component of noise covariance should be reduced when feedback from decision areas – or areas mediating feedback signals – is blocked from arriving to the recorded sensory area.

### Inferring variable internal beliefs from sensory responses

We have shown that internal beliefs about the stimulus induce corresponding structure in the correlated variability of sensory neurons’ responses (Figure 7a). Conversely, this means that the statistical structure in sensory responses can be used to infer properties of those beliefs.

**Figure 7.**
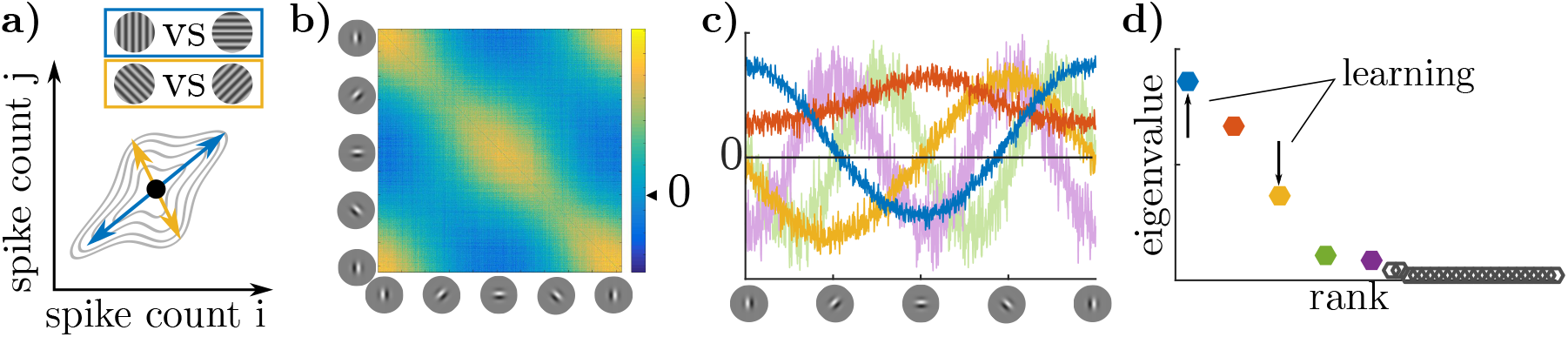
Inferring internal beliefs. **a)** Trial-to-trial fluctuations in the posterior beliefs about **x** imply trial-to-trial variability in the mean responses representing that posterior. Each such ‘belief’ yields increased correlations in a different direction in **r**. The model in (b-d) has uncertainty in each trial about whether the current task is a vertical-horizontal orientation discrimination (task 1, blue) or an oblique discrimination (task 2, yellow). **b)** Correlation structure of simulated sensory responses during discrimination task. Neurons are sorted by their preferred orientation (based on (Haefner et al., 2016)). **c)** Eigenvectors of correlation matrix (principal components) plotted as a function of neurons’ preferred orientation. The blue vector corresponds to fluctuations in the belief that either a vertical or horizontal grating is present (task 1), and the yellow corresponds to fluctuations in the belief that an obliquely-oriented grating is present (task 2). See Methods for other colours. **d)** Corresponding eigenvalues color-coded as in (c). Our results on variable beliefs (*π*) predict an increase over learning in the eigenvalue corresponding to fluctuations in belief for the correct task, while our results on filtering noise predict only a relative increase in the task-relevant eigenvalue compared with variance in other tasks’ directions (e.g. if both blue and yellow decrease, but yellow more so).

In order to demonstrate the usefulness of this approach, we used it to infer the structure of an existing model for which we know the ground truth (Haefner et al., 2016). The model discriminated either between a vertical and a horizontal grating (cardinal context), or between a –45deg and +45deg grating (oblique context). The model was given an unreliable (80/20) cue as to the correct context before each trial, and thus had uncertainty about the exact context. The model simulates the responses of a population of primary visual cortex neurons with oriented receptive fields that perform sampling-based inference over image features. Since the relevant stimulus dimension for this task is orientation, we sorted the neurons by preferred orientation. The resulting noise correlation matrix – computed for *zero-signal trials* – has a characteristic structure in qualitative agreement with empirical observations (Figure 7b) (Bondy et al., 2018).

We found that the simulated neural responses had five significant principal components (PCs) when the true context was cardinal discrimination (Figure 7c-d). Knowing the preferred orientation of each neuron allows us to interpret the PCs as directions of variation in the model’s belief about the current orientation. For instance, the elements of the first PC (blue in Figure 7c) are largest for neurons preferring vertical and negative for those preferring horizontal orientation, indicating that there is trial-to-trial variability in the model’s internal belief about whether “there is a vertical grating and not a horizontal grating” – or vice versa – in the stimulus, corresponding to the **f′**–axis of the cardinal task. Analogously, one can interpret the third PC (yellow in Figure 7c-d) as corresponding to the belief that a +45° grating is being presented, but not a –45° grating, or vice versa. This is the **f′**-axis for the wrong (oblique) task context, reflecting the fact that the model maintained some uncertainty about which was the correct task in a given trial. The remaining PCs in Figure 7c-d correspond to task-independent variability (see Supplemental Figure S3).

Maintaining uncertainty about the task itself is the optimal strategy from the subject’s perspective given their imperfect knowledge of the world. When compared to perfect knowledge of context, it decreases behavioral performance. Behavioral performance is optimal only when the internal model learned by the subject exactly corresponds to the experimenter-defined one – an ideal which subjects should approach over the course of learning. An empirical prediction, therefore, is that eigenvalues corresponding to the correct task-defined stimulus dimension will increase with learning, while eigenvalues representing other tasks should decrease. Furthermore, the shape of the task-relevant eigenvectors should be predictive of psychophysical task-strategy. Importantly, they constitute a richer, higher-dimensional, characterization of a subject’s decision strategy than psychophysical kernels or CPs (Nienborg and Cumming, 2007) (Figure 7c).

## Discussion

We derived a novel analytical link between the two dominant frameworks for modeling sensory perception: probabilistic inference and neural population coding. Under the assumption that neural responses represent posterior beliefs, we showed how trial-to-trial variability in those beliefs induces empirically observable covariability in neural responses. Exploiting a fundamental self-consistency relationship underlying Bayesian learning, we were able to make specific predictions for the nature of neural and behavioral correlations in classic discrimination tasks with almost no assumptions about how beliefs are encoded in neural responses. Re-examining existing data we found evidence for these predictions, both supporting the hypothesis that neurons encode posterior beliefs and providing a novel explanation for previously puzzling empirical observations. Finally, we illustrated how measurements of neural responses can in principle be used to infer a subjects internal beliefs in the context of a task.

### Feedback and correlations

Our results directly address several debates in the field on the nature of feedback to sensory populations. First, they provide a rationale for the apparent ‘contamination’ of sensory responses by top-down decision signals (Nienborg and Cumming, 2009; Wimmer et al., 2015; Ecker et al., 2016; Rabinowitz et al., 2015; Bondy et al., 2018; Haimerl et al., 2019): top-down signals communicate task-relevant expectations, not reflecting the decision *per se* but integrating information about the outside world (Nienborg and Roelfsema, 2015). Second, this feedback may be dynamic, reflecting the subject’s growing confidence within a trial and inducing choice probabilities that are the result of both feedforward and (growing) feedback components (Nienborg and Cumming, 2009; 2014; Wimmer et al., 2015; Haefner et al., 2016). Third, these feedback signals also introduce correlated sensory variability that is information-limiting (Moreno-Bote et al., 2014) in tasks in which integrating some information may not be warranted, e.g. because individual stimuli and trials are temporally uncorrelated.

We identified three distinct mechanisms by which correlated variability arises in a Bayesian inference framework. The first is neural variability in the encoding of a fixed posterior. This type of variability has previously been studied especially in neural sampling codes (Hoyer and Hyvärinen, 2003; Orbán et al., 2016; Echeveste et al., 2019; Bányai et al., 2019; Bányai and Orbán, 2019). Instead, we study variability in the posterior itself, which arises due to both task-dependent and task-independent mechanisms. The second mechanism is variability in task-relevant categorical belief (*π*), projected back to sensory populations during each trial. Under conditions consistent with threshold psychophysics, we showed that variable categorical beliefs induce commensurate choice probabilities and neural covariability in approximately the **f′**–direction assuming the subject learns optimal statistical dependencies. This holds for general distributional codes if noise is negligible, and for a newly-identified class of Linear Distributional Codes (LDCs) in the case of non-negligible noise. The third source of variability in neural responses is due to task-independent noise that interacts with a task-dependent prior. Although not solved analytically, we found in simulation that the task-dependent component of this variability likewise implies increased differential correlations after learning, though not necessarily increased differential covariance. The latter two mechanisms act through feedback: in one case there is dynamic feedback of a particular belief *π*, and in the other case there is task-dependent (but belief-independent) feedback that sets a static prior each trial, then interacts with noise in the likelihood, analogous to models of “state-dependent” recurrent dynamics (Huang et al., 2019; Doiron et al., 2016; Ramalingam et al., 2013).

Of these two mechanisms, empirical data on choice probabilities suggests that variability in belief (*π*) may dominate in many existing studies. Choice probabilities could in theory arise from a combination of three mechanisms: (i) feedforward causal effects of sensory neurons on behavior (Shadlen et al., 1996; Haefner et al., 2013; Pitkow et al., 2015), (ii) across-trial autocorrelation of both behavior and neural activity acting independently (Lueckmann et al., 2018), or (iii) feedback of belief or choice within a trial (Nienborg and Cumming, 2009; Wimmer et al., 2015; Haefner et al., 2016). Our analysis of variability in *π* is compatible with (iii), while variable likelihoods would be compatible with (i). Experimental work has suggested that both (i) and (ii) are insufficient to account for a large fraction of choice probability (Nienborg and Cumming, 2009; Wimmer et al., 2015; Lueckmann et al., 2018). Interpreted in our framework, this suggests that feedback of variable beliefs has a greater overall effect on the task-dependent statistics of neural activity than variable likelihoods, at least in those tasks and brain areas.

Our results suggest that at least some of measured “differential” covariance may be usefully understood as near-optimal feedback from internal belief states or as the interaction between task-independent noise and a task-specific prior. In neither case is information necessarily more limited as the result of learning. In the first case, while feedback of belief (*π*) biases the sensory population, that bias may be accounted for by downstream areas (Kohn et al., 2016; Chicharro et al., 2017). In principle, these variable belief states could *add* information to the sensory representation if they are *true* (Lange and Haefner, 2017). In the second case, the noise in the **f′** direction *does* limit information, but to the same extent as before learning; there is not necessarily *further* reduction of information by “shaping” the noise with a task-specific prior. For a fixed population size, it is covariance in the **f′** direction, not correlation, that ultimately affects information.

### Posterior Coding

Our focus on firing rates and spike count covariance is motivated by connections to rate-based encoding and decoding theory. We do not assume that they are the sole carrier of information about the underlying posterior p_b_(**x**|…), but simply statistics of a larger spatio-temporal space of neural activity, **r** (Dayan and Abbott, 2001). For many distributional codes, firing rates are only a summary statistic, but they nonetheless provide a window into the underlying distributional representation.

Probabilistic Population Codes (PPCs) have been instrumental for the field’s understanding of the neural basis of inference in perceptual decision-making. However, they are typically studied in a purely feedforward setting assuming a representation of the likelihood, not posterior (Ma et al., 2006; Beck et al., 2008). In contrast, Tajima et al. (2016) modeled a PPC encoding the posterior and found that categorical priors bias neural responses in the **f′** direction, consistent with our results (Tajima et al., 2016).

The assumption that sensory responses represent posterior beliefs through a general encoding scheme agrees with empirical findings about the top-down influence of experience and beliefs on sensory responses (von der Heydt et al., 1984; Lee and Mumford, 2003; Nienborg and Cumming, 2014). It also relates to a large literature on association learning and visual imagery (reviewed in (Albright, 2012)). In particular, the idea of ‘perceptual equivalence’ (Finke, 1980) reflects our starting point that the very same posterior belief (and hence the same percept) can be the result of different combinations of sensory inputs and prior expectations. In a discrimination task, for instance, there are three distinct associations inducing correlations. First, showing the same input many times induces positive correlations between sensory neurons responding to the same input. Second, presenting only one of two possible inputs induces negative correlations between neurons responding to different inputs. Third, keeping the input constant within a trial induces positive auto-correlations. All three associations are directly reflected in the predicted (Figure 7b, Haefner et al. (2016)), and empirically observed neural responses (Bondy et al., 2018; Lueckmann et al., 2018).

Our derivations implicitly assumed that the feedforward encoding of sensory information, i.e. the likelihood p(**E**|**x**), remains unchanged between the compared conditions. This is well-justified for lower sensory areas in adult subjects (Hensch, 2005), or when task contexts are switched on a trial-by-trial basis (Cohen and Newsome, 2008). However, it is not necessarily true for higher cortices (Li and DiCarlo, 2008), especially when the conditions being compared are separated by long periods of task (re)training (Bondy et al., 2018). In those cases, changing sensory statistics may lead to changes in the feedforward encoding, and hence the nature of the represented variable **x** (Ganguli and Simoncelli, 2014; Wei and Stocker, 2015).

### Outlook

We introduced a general notation for distributional codes, 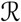, that encompasses nearly all previously proposed distributional codes. Thinking of distributional codes in this way – as a map from an implicit space p_b_(**x**) to observable neural responses p(**r**) – is reminiscent of early work on distributional codes (Zemel et al., 1998), and emphasizes the convenience of computation, manipulation, and decoding of p_b_(**x**|…) from r rather than its spatial or temporal allocation of information *per se* (Fiser et al., 2010; Pouget et al., 2013; Gershman and Beck, 2016). Our results leverage this generality and show that properties of Bayesian computation might be identified in neural populations without strong commitments to its algorithmic implementation. Rather than assuming an approximate inference algorithm (e.g. sampling) then deriving predictions for neural data, future work might productively work in the reverse direction, asking what class of generative models (**x**) and encodings 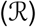 are consistent with some data. As an example of this approach, we observe that the results of Berkes et al. (2011) are consistent with any LDC, since LDCs have the property that the average of encoded distributions equals the encoding of the average distribution, exactly as the authors reported (Berkes et al., 2011).

Distinguishing between linear and nonlinear distributional codes is complementary to the much-debated distinction between parametric and sampling-based codes. LDCs include both sampling codes where samples are linearly related to firing rate (Hoyer and Hyvärinen, 2003; Buesing et al., 2011; Pecevski et al., 2011; Savin and Denève, 2014; Haefner et al., 2016; Shivkumar et al., 2018) as well as parametric codes where firing rates are proportional to expected statistics of the distribution (Anderson and Van Essen, 1994; Zemel et al., 1998; Sahani and Dayan, 2003; Vertes and Sahani, 2018). Examples of distributional codes that are *not* LDCs include sampling codes with nonlinear embeddings of the samples in r (Aitchson and Lengyel, 2016; Orbán et al., 2016; Echeveste et al., 2019) and parametric codes in which the *natural parameters* of an exponential family are encoded (Ma et al., 2006; Beck et al., 2008; 2013; Raju and Pitkow, 2016).

Our results provide a normative justification for decision-related feedback that is aligned with *vf′*. In the context of our theory, there are three possible deviations from our assumptions that can account for empirical results of a less-than-perfect alignment (Ni et al., 2018) – each of them empirically testable. First, it is plausible that only a subset of sensory neurons represent the posterior, while others represent information about necessary ‘ingredients’ (likelihood, prior), or carry out other auxiliary functions (Pecevski et al., 2011; Aitchson and Lengyel, 2016). Our predictions are most likely to hold among layer 2/3 pyramidal cells, which are generally thought to encode the *output* of cortical computation in a given area, i.e. the posterior in our framework (Felleman and Van Essen, 1991). Second, subjects may not learn the task *exactly* implying a difference between the experimenter-defined task and the subject’s ‘subjective” **f′** direction for which our predictions apply. This explanation could be verified using psychophysical reverse correlation identifying the subject’s “subjective” **f′** direction from behavioral data. Finally, some misalignment between **f′** and decision-related feedback may be indicative of significant task-independent noise in the presence of a nonlinear distributional code, which could be tested by manipulating the amount of external noise in the stimulus.

Much research has gone into inferring latent variables that contribute to the responses of neural responses (Cunningham and Yu, 2014; Archer et al., 2014; Kobak et al., 2016). Our predictions suggest that at least some of these latent variables can usefully be characterized as internal beliefs about sensory variables. We showed in simulation that the influence of each latent variable on recorded sensory neurons can be interpreted in the stimulus space using knowledge of the stimulus-dependence of each neuron’s tuning function (Figure 7). Our results are complementary to *behavioral* methods to infer the shape of a subject’s prior (Houlsby et al., 2013), but have the advantage that the amount of information that can be collected in neurophysiology experiments far exceeds that in psychophysical studies allowing for richer characterization of the subject’s internal model (Ruff et al., 2018).

The detail with which internal beliefs can be recovered from the statistical structure in neurophysiological recordings is limited by both experimental and theoretical techniques. While much current research is aimed at developing those techniques and at characterizing the latent structure in the resulting recordings, how to make sense of the observed structures is less clear. Our work suggests a way to interpret this structure, and makes predictions about how it should change with task context and learning.

## Methods

### Optimal task-induced sensory expectations

Following previous work (Olshausen and Field, 1996; Lee and Mumford, 2003; Kersten et al., 2004; Fiser et al., 2010), we assume that the brain has learned an implicit hierarchical generative model of its sensory inputs, p_b_(**E**|**x**), in which perception corresponds to inference of latent variables, **x**, conditioned on those inputs. The subscripted distributions p_b_(·) and p_e_(·) refer to the brain’s internal model and the experimenter’s “ground truth” model, respectively (Figure 4a).

In the classic two-alternative forced-choice (2AFC) paradigm, the experimenter parameterizes the stimulus with a scalar variable *s* and defines category boundary which we will arbitrarily denote *s* = 0. If there is no external noise, the scalar *s* is mapped to stimuli by some function **E**(*s*), for instance by rendering grating images at a particular orientation. In the case of noise, below, we consider more general stimulus distributions p_e_(**E**|*s*).

We assume that the brain does not have an explicit representation of *s* but must form an internal estimate of the category each trial, *Ĉ*, based on the variables represented by sensory areas, **x** (Shivkumar et al., 2018). From the “ground truth” model perspective, stimuli directly elicit perceptual inferences – this is why we include p_e_(**x**|**E**) as part of the experimenter’s model. In the brain’s internal model, on the other hand, the stimulus is assumed to have been generated by causes x, which are, in turn, *jointly* related to *Ĉ*. These models imply the following conditional independence relations (Figure 4a+b):

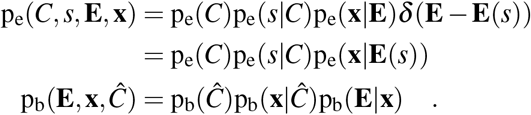

We assume the brain learns the joint distribution p_b_(**x**, *Ĉ*) that maximizes reward, or equivalently that best matches the ground-truth distribution p_e_(*C*,**x**) in expectation (Figure 4a). This entails a conditional distribution “decoding” *Ĉ* from **x** of the form

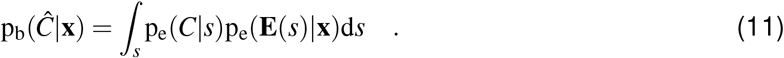

We next derive the reciprocal influence of *Ĉ* on **x** (equation (2) in the main text) by applying Bayes’ rule to equation (11):

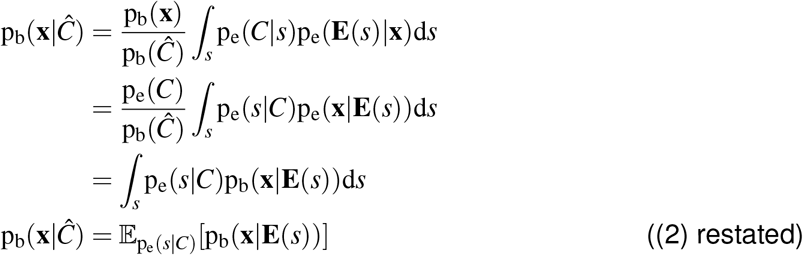

The substitution of p_b_ for p_e_ in the third line follows from the fact that, even from the perspective of an external observer, p_e_(**x**|*s*) is the inference made *by the brain* about **x** induced by the stimulus **E**(*s*). Hence, p_e_(**x**|*s*) is equivalent to p_b_(**x**|**E**(*s*)). The fractions p_e_(*C*)/p_b_(*Ĉ*) and p_b_(**x**)/p_e_(**x**) become one, assuming that the subject learns the correct categorical prior on C and a consistent internal model. We note that this distribution can be learned even if *s* is not directly observable by the brain, since its model has access to the true category labels if subjects are informed of the correct answer each trial, as well as to each individual posterior p_b_(**x**|*s*), as this is what we assume is represented by the sensory area. See the Supplemental Text for further discussion of this expression.

As described in the main text we marginalize over the subject’s belief in the category, *π* = p_b_(*Ĉ* = 1), to get an expression for expectations on **x** given the belief (equation (3)). Unlike *Ĉ*, *π* is not a random variable in the generative model but the *parameter* defining the subject’s belief about the binary variable *Ĉ*. The resulting posterior on **x**, abbreviated in equation (4), is given by

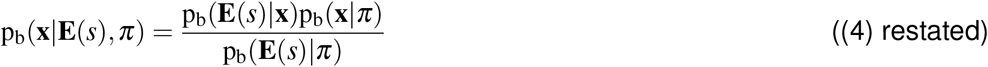

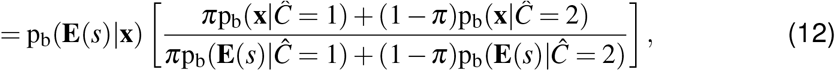

We assume that the category boundary *s* = 0 is itself equally likely to occur conditioned on each category (usually true by definition), but note that this is *not* a requirement that the categories are *a priori* equally likely. This simplifies equation (12) when conditioning on *s* = 0:

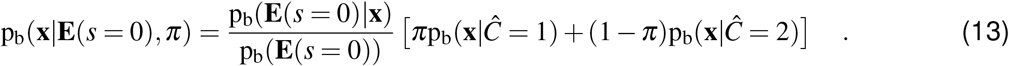

### Proof of approximate proportionality of derivatives of the posterior (5)

Our first main result is the approximate proportionality in (5), restated here:

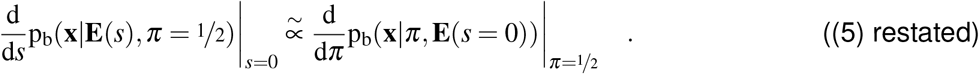

We use *π* = ½ to denote the true prior over categories, which is often 50/50 but our results hold for biased p_e_(*C*) as well.

Since *s* = 0 is fixed in the right-hand-side of (5), the total derivative with respect to *π* equals its partial derivative, assuming that there are no *additional* internal variables that are dependent on both **x** and *π*. In the left-hand-side of (5), the total derivative with respect to *s* includes two terms, one due to the direct effect of *s* on the posterior, and the other due to the mean dependence of *π* on s, since changes in *s* elicit changes in the subject’s beliefs:

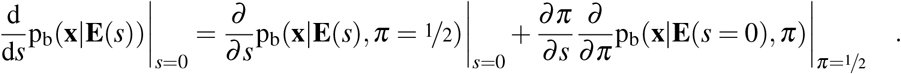

Below, we will replace p_b_(**x**|**E**(*s*), *π* = ½) with p_b_(**x**|**E**(*s*)) to reduce notational clutter since *π* = ½ corresponds to marginalizing over categories with the true prior. The second partial derivative term in the previous equation is equal to the right-hand-side of (5), scaled by *∂_π_/∂_s_*, and hence does not affect the overall proportionality in (5). To prove the approximate proportionality in (5), we therefore need only prove proportionality in the partial derivatives:

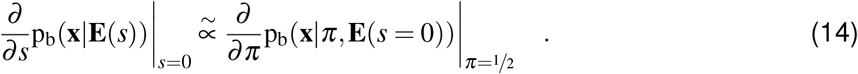

Using a small Δ*s* finite-difference approximation, we rewrite t the left-hand-side of (14) as

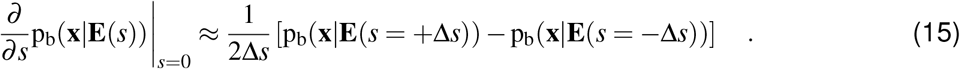

While this is an approximation to the “true” derivative, it is usually a good one based on theoretical reasons (range of *s* small in the threshold regime of psychophysical tasks) and empirical observations (Bondy et al., 2018).

Next, consider the right-hand-side of (14) using the expression for the posterior conditioned on *s* = 0 (equation (13)). The partial derivative of this posterior with respect to the belief *π* is

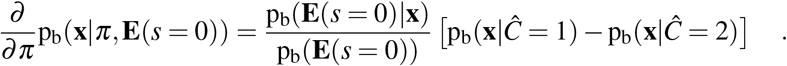

Applying the self-consistency constraint implied by learning (i.e. substituting in equation (2) to the terms inside the brackets), this becomes

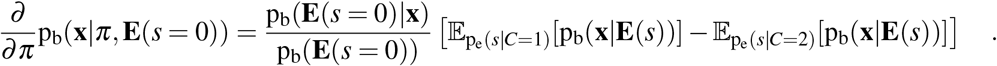

Re-arranging terms, we arrive at

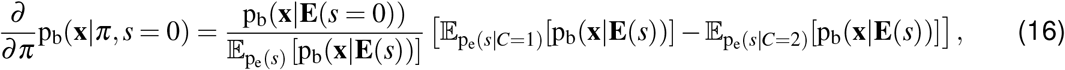

where we have used the identity 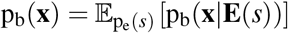 to write the denominator of the fraction outside the brackets as expectations over s. This identity is valid because we assumed subjects have completely learned the task, so the *self-consistency* rule holds that the prior p_b_(**x**) equals the average posterior seen in the task (Dayan and Abbott, 2001).

Having re-arranged terms, we must now establish conditions under which (15) and (16) are proportional. While they appear similar by inspection, they are not proportional in general because so far we have placed no restrictions on the experimenter’s distribution of stimuli p_e_(*s*). We therefore next consider the special case of sub-threshold tasks. One way to formalize this mathematically is by taking the limit of (16) as p_e_(*s*) approaches a Dirac delta around *s* = 0, as this appears to result in agreement between the individual terms of (16) and (15). However, in this limit (16) itself goes to zero (indeed, it should be expected that beliefs are irrelevant in a task that has zero variation in stimuli).

This suggests an approximate solution by breaking the problem into two limiting processes: one in which the distribution of stimuli within each category concentrates on some ±Δ*s*, and a second in which Δ*s* gets small (but does not reach zero). Supplemental Figure S1 visualizes these two steps. To realize the first limit, we set

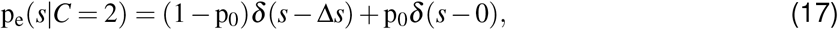

and likewise for *C* = 1 and −Δ*s*. We include the *δ*(*s* – 0) term to ensure that zero-signal stimuli are always included with probability p_0_, otherwise evaluating (16) at *s* = 0 would not be possible in practice. Marginalizing over categories, the full distribution of stimuli becomes

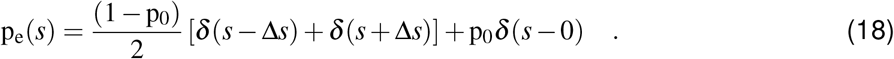

Substituting equations (17) and (18) into (16) simplifies the expectations. First, the terms inside the brackets in (16) goes to

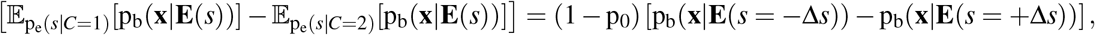

which matches the corresponding term in (15) to the extent that Δ*s* is small enough to approximate the derivative 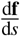. Thus, the extent to which (16) is proportional to (15) depends only on the extent to which the first term in the right-hand-side of (16) is constant, or equivalently whether p_b_(**x**|**E**(*s* = 0)) approximately equals 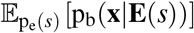. Considering the special case of stimulus distributions given in (17) and (18), this near-equality condition holds as the probability of true zero-signal stimuli (p0) grows, or as the category differences (Δ*s*) shrink: an approximation to sub-threshold psychophysics conditions.

Taken together, this establishes the approximate proportionality in (14), which in turn concludes the proof of (5), in the special case of sub-threshold psychophysics. See the Supplemental Text for further discussion of the applicability and interpretation of these limits.

### Encoding the posterior in neural responses

Our above derivations considered perturbations of an approximate Bayesian observer’s posterior over their internal variables, p_b_(**x**|**E**(*s*), *π*). We next link these computational-level changes in the posterior to predictions for observable changes in neural firing rate. “Posterior coding” hypothesizes that the (possibly high-dimensional) posterior p_b_(**x**|**E**(*s*),*π*) is encoded in the spiking pattern of a population of neurons over some time window. We do not restrict the space of neural responses **r** to total spike counts or average spike rates, but instead consider r on a single trial to live in a high-dimensional “spatiotemporal” space, i.e. an *N* × *B* array of spike counts for all N neurons in a population resolved into B fine-timescale bins (Dayan and Abbott, 2001). That is, 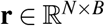, where **r**_ib_ is the spike count of neuron *i* at time *b*. This definition subsumes both “spatial” and “temporal” codes, a distinction that lies at the center of some debates over the neural representation of distributions (Fiser et al., 2010; Pouget et al., 2013; Gershman and Beck, 2016).

We define distributional codes of the *posterior* as any encoding scheme 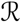 where the posterior distribution on **x** is sufficient to determine the neural response distribution over the range of possible stimuli^3^. Formally, we say

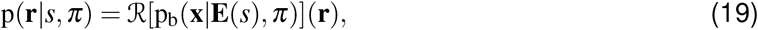

where 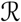 is a higher-order function that maps from distributions over **x** to distributions over **r**. (One may equivalently think of 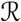 either as a deterministic higher-order map as we have written here, or as a stochastic map from distributions on **x** directly to neural activity patterns r.) Our only restrictions on **x** and 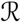 are that p_b_(**x**|…) must be sufficiently wide, and 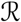 must be sufficiently smooth over the relevant range of stimulus values, so that the derivatives and linear approximations throughout are valid. A second restriction on **x** and 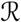 is that the dominant effect of *s* on **r** must be in the mean firing rates rather than their higher-order moments of **r**. While this is a theoretically complex condition to meet involving interactions between *s*, **x**, and 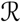, it is easily verified empirically in a given experimental context: if changes to *s* primarily influence the mean spike count, it is irrelevant whether these changes coded for the mean, variance, or higher-order moments of p_b_(**x**|…). If the space of **r** is the full “spatiotemporal” space of neural activity patterns, this definition encompasses all previously proposed parametric (Beck et al., 2013; Raju and Pitkow, 2016; Tajima et al., 2016; Vertes and Sahani, 2018), and sampling-based (Hoyer and Hyvärinen, 2003; Buesing et al., 2011; Savin and Denève, 2014; Orbán et al., 2016; Haefner et al., 2016; Aitchson and Lengyel, 2016) encoding schemes as special cases, among others. However, it excludes sub-populations of neurons in which only the likelihood or prior, but not the posterior, is encoded (Ma et al., 2006; Beck et al., 2008; Walker et al., 2019).

### Tuning curves as statistics of encoded distributions

The total spike count of neuron *i* in terms of **r** is a function of **r** that sums responses over time bins:

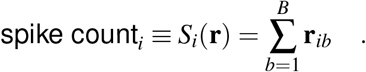

In an encoding model defined as in equation (19), each neuron’s tuning curve is thus defined by the expectation of *S_i_* at each value of the stimulus *s*:

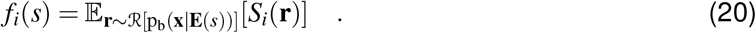

The *slope* of this tuning curve, 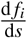, is given by the chain rule:

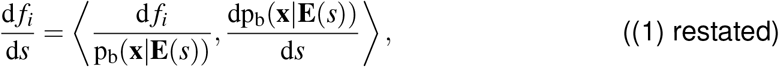

where the inner product is taken between two functions, since derivatives were taken with respect to the distribution p_b_(**x**|**E**(*s*), *π*). Equation (1) shows how we use smoothness and linearization assumptions to decouple our analysis of changes in posteriors (e.g. dp_b_/d*s*) from their effect on mean firing rates under arbitrary distributional encodings (e.g. d*f_i_*/dp_b_). The proportionality between dp_b_/d*s* due to changing stimuli and dp_b_/d*π* due to feedback of beliefs (equation (5)) implies an analogous proportionality in neural responses:

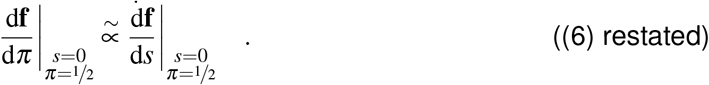

### Implication for top-down component of choice probability

We assume the subject’s choice is based on their posterior belief in the stimulus category, i.e. value of *π*. Conditioning neural responses on choice is then equivalent to conditioning on the sign of *π* – ½ (if there is an additional stage of randomness between belief *π* and behavioral choice, what follows will remain true up to a proportionality, (Chicharro et al., 2017)).

Let CTA_*i*_ be the “choice triggered average” of neuron ¡, defined as the difference in mean response to choice 1 and choice 2. To isolate top-down effects, consider the noiseless case where neural responses depend exclusively on *s* (which is fixed) and *π* (which is varying). We then write CTA as the difference in expected neural response between the *π* > ½ and *π* < ½ cases:

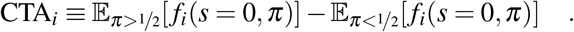

For small variability in *π*, this can be approximated linearly:

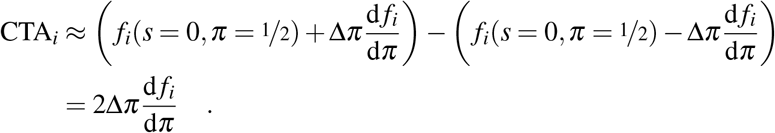

Substituting in the proportionality 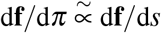 (6), it follows that 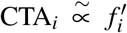. Dividing both sides of this proportionality by the standard deviation of the neuron’s response, *σ*_*j*_, and incorporatig the fact that 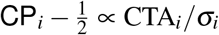 (Haefner et al., 2013; Pitkow et al., 2015), we arrive at the following equation for the *top-down* component of choice probability after learning:

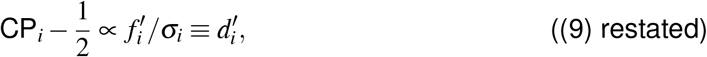

where *d′* is the “d-prime” sensitivity measure from signal detection theory (Green and Swets, 1966).

### Implication for task-dependence of noise covariance

Consider any scalar variable *a* that linearly shifts neural responses in an arbitrary direction u, above and beyond all of the other factors influencing the population (denoted “…”):

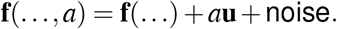

When *a* varies from trial to trial, it adds a rank-1 component to the covariance matrix:

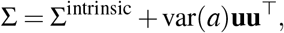

where Σ^intrinsic^ is the covariance due to all other factors, i.e. due to neural noise and variability in any of the terms in “…”.

It follows that *variability* in the posterior along dp_b_/d*s* manifest as covariability among neurons in the **f′f′**^T^ direction (Lange and Haefner, 2017). The noise covariance structure due to var(*π*) is predicted to be

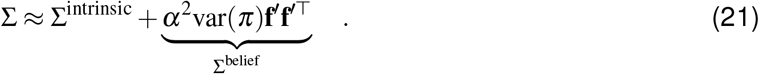

Σ^intrinsic^ may be thought of as neural noise above and beyond variability in belief. Σ^belief^ is the rank-one component in the **f′f′**^T^ direction due to feedback of variable beliefs, and *α* is the proportionality constant from (5).

We call two tasks ‘comparable’ when they agree both in the magnitude of their top-down effects (*α*^2^var(*π*)) and in their intrinsic response covariance (Σ^intrinsic^), as can reasonably be expected, for instance, in rotationally symmetric coarse discrimination tasks where all that changes between the tasks is the orientation (Bondy et al., 2018) or motion direction (Cohen and Newsome, 2008) of the discrimination boundary while the zero-signal stimulus stays the same. In that case subtracting the covariance matrices from each task yields the following prediction (Figure 6b):

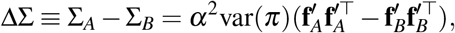

having cancelled out the task-independent term Σ^intrinsic^.

Note that two fine discrimination tasks (e.g. orientation discrimination around the vertical and the horizontal axes, respectively) are not necessarily ‘comparable’ since the two tasks differ in their zero-signal stimulus (a vertical and a horizontal grating, respectively), which may yield different baseline covariability, Σ^intrinsic^.

### Inferring the internal model

Complex tasks (e.g. those switching between different contexts), or incomplete learning (e.g. uncertainty about fixed task parameters), will often induce variability in multiple internal beliefs about the stimulus. Assuming that this variability is independent between the beliefs, we can write the observed covariance as Σ ≈ Σ^0^ + ∑_k_*λ*^(*k*)^**u**^(*k*)^**u**^(*k*)T^. Here, each vector **u**^(*k*)^ corresponds to the change in the population response corresponding to a change in internal belief *k*. The coefficients *λ*^(k)^ are proportional to the variance of the trial-to-trial variability in belief *k*, as in var(*π*) above, and Σ^0^ represents all task-independent covariance.

The model in our proof-of-concept simulations has been described previously (Haefner et al., 2016). In brief, it performs inference by neural sampling in a linear sparse-coding model (Olshausen and Field, 1996; Hoyer and Hyvärinen, 2003; Fiser et al., 2010). The prior is derived from an orientation discrimination task with two contexts – oblique orientations and cardinal orientations - that is modeled on an analog direction discrimination task (Cohen and Newsome, 2008). We simulated the responses of 1024 V1 neurons whose receptive fields uniformly tiled the orientation space. Each neuron’s response corresponds a set of samples from the posterior distribution over the intensity of its receptive field in the input image. We simulated zero-signal trials by presenting white noise images to the model. The eigenvectors not described in the main text correspond to stimulus-driven covariability, plotted in Figure S3 for comparison.

### Task-independent variability in the posterior

We consider three potential sources of task-independent noise in posteriors: first, there are additional “high level” variables in *I* that may be probabilistically related to **x** but are not task-relevant. Just as variability in *π* induces variability in p_b_(**x**|**E**(*s*), *π*), variability in these other internal states may induce variability in the posterior. Second, there may be measurement noise in the observation of E or noise in the neurons afferent to those representing **x**. Third, the stimulus itself may be stochastic by design, drawn according to some p_e_(**E**|*s*). We model these sources of variability by three types of noise, *ε* = {*ε*_**I**_, *ε*_*L*_, *ε*_**E**_} corresponding to “internal state” noise, “likelihood” noise, and stimulus noise respectively. We assume that the all noise sources are unaffected by task learning or task context and are independent of both *s* and *π*.

By approximating the joint effect of *π* and *ε*_I_ on the density of **x** as multiplicative, the full posterior decomposes as follows:

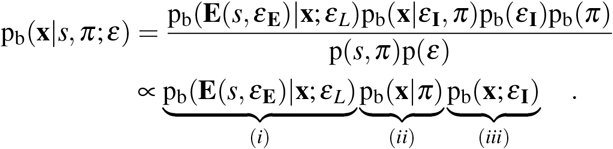

The first term (*i*) is the “noisy likelihood” conditioned on the noisy stimulus **E**(*s*, *ε*_E_). The second term (*ii*) is the task-dependent component of the prior studied above. The third term (*iii*) captures the influence due to other internal variables besides *π*.

The two noise terms, (*i*) and (*iii*), may be combined into a single term. With some slight abuse of notation, we can replace p_b_(**E**(*s*,*ε*_**E**_)|**x**;*ε*_L_) with p_b_(*s*|**x**;*ε*_L_,*ε*_**E**_) so that the ε terms appear together. Combining terms, one can thus interpret both (*iii*) and (*i*) as noise in the likelihood, despite one being feed-back and the other being feed-forward:

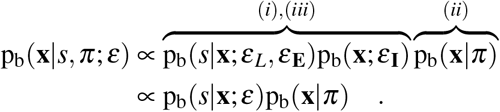

This motivates our discussion only of “noisy likelihoods” in the main text – it implicitly includes stimulus noise, feedforward noise, and noise due to variable internal states besides *π*.

### Variable beliefs in the presence of noise

Analogous to equation (2) in the main text, learning the task in the the presence of noise implies learning a prior that is equal to the average of (noisy) posteriors seen in the task:

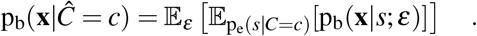

Paralleling the deriviation of (3), this implies a prior conditioned on the graded belief *π* of the form

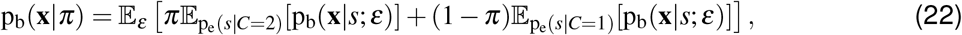

which is identical to (3), but with the average posteriors further “blurred” by the noise.

The expected spike count of neuron *i*, denoted *f_i_*, previously contained only an expectation over neural responses **r**; now we simply add an outer expectation over *ε*:

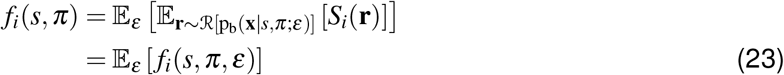

where *S_i_*(**r**) is again simply counts the spikes of neuron *i*. The second line defines a new three-argument function *f_i_*(*s*, *π*, *ε*) which is the expected spike count of neuron *i* for fixed *s*, *π*, and *ε*.

We again consider the case of zero-signal stimuli and the relationship between 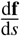 and 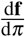. As before, the population’s sensitivity to the stimulus, 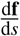, is approximated by the average difference between **f**(+Δ*s*) and **f**(–Δ*s*) (analogous to equation (15) which estimated 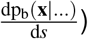:

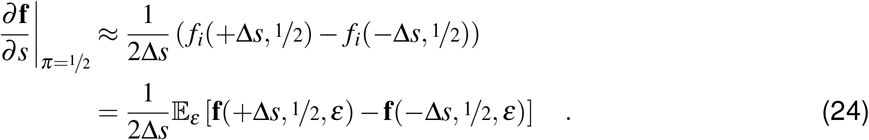

Note that by reparameterizing p_e_(**E**|*s*) as the deterministic function **E**(*s*, *ε*_**E**_), we are able to pass the derivative with respect to *s* through expectations over *ε*, as in the “reparameterization trick” (Rezende et al., 2014).

We again apply the chain rule to express the population’s sensitivity to beliefs *π* in the presence of noise as an expectation over an inner product:

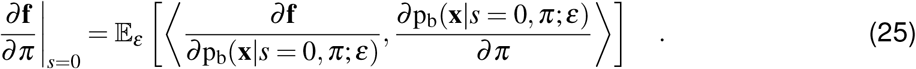

From (22), we have

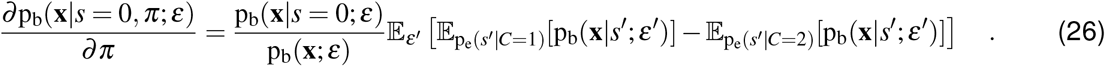

Following our proof of (5), we again assume the case of narrow stimulus distributions (equation (17)) in the sub-threshold regime (so Δ*s* is small). The outer expectation over *ε* in (25) only affects the term 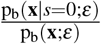 in (26), and this term again becomes negligible in the sub-threshold limit. The inner expectation over *ε′* remains, however.

Comparing (24) with (25)-(26), the effect of noise becomes apparent: while 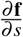 has the form of an *expectation of the difference* of **f** evaluated across noise values, 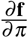 has the form of **f** evaluated on the *difference of expectations*. Unlike in the noiseless case, these are no longer proportional in general.

However, we observe that proportionality between (24) and (25) still holds for a restricted class of distributional encoding schemes 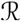, namely those distributional codes for which *firing rates are linear in mixtures of distributions*. Let p_3_ (**x**) be a mixture of two distributions, *α*p_1_(**x**) + (1 – *α*)p_2_(**x**), 0 ≤ *α* ≤ 1. Formally, we define “Linear Distributional Codes” (LDCs) as all codes for which the following holds for all p_1_ and p_2_:

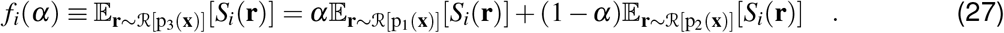

LDCs have the property that the expectation over *ε* pass through the function **f**(). Combined with (24)-(26), this implies that in cases with significant task-independent noise, only linear distributional codes will have the property that 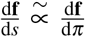, and hence make all the same predictions for data described in the main text, such as the emergence of both differential correlations and a top-down component of choice probabilities proportional to neural sensitivities over learning.

### Interactions between task-independent noise and task-dependent priors

Throughout this section, we will fix *s* = 0 and *π* = ½ to isolate the effects of ε in “zero-signal” conditions. We will also assume that **x** is discrete so that we can use finite-length vectors of probability mass rather than probability density functions, but this is only for intuition and notational convenience.

Above, we used the chain rule of derivatives to write neurons’ sensitivity to various factors in terms of their sensitivity to the posterior density, d**f**/dp_b_(**x**|…). To a first approximation, the same trick can be applied to write the *covariance* of neural responses in terms of their sensitivity to p_b_(**x**|…) and the *covariance* in the posterior mass itself due to task-independent noise (*ε*):

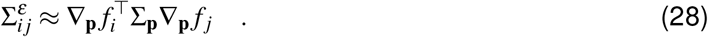

The inner term, Σ_p_, is the *covariance of the elements of the posterior* p_b_(**x**|…) *at pairs of points* **x**_1_, **x**_2_ *due to ε* (see Supplemental Text for further discussion of this term). The term ∇_p_*f_i_* is the gradient of neuron *i*’s firing rate with respect to the elements of p_b_(**x**|…).

Recall that the noisy posterior, p_b_(**x**|*s*, *π*; *ε*), can be written with all noise terms in the likelihood, i.e. p_b_(x|*π*)p_b_(s|x;ε) (up to constants). Because of this, the prior may be pulled out of Σ_p_ as follows (we drop *π* = ½ here to reduce clutter):

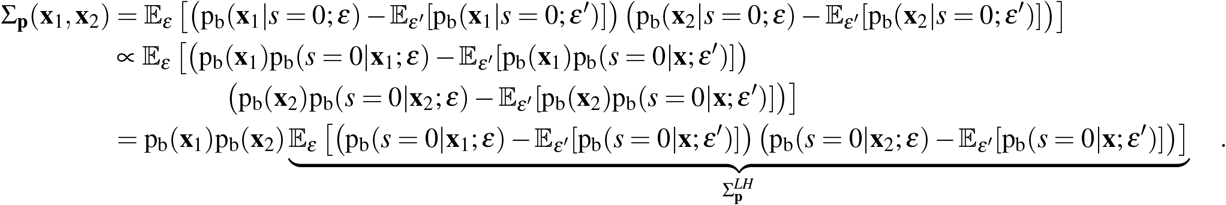

In the second line, we absorbed p_b_(*s* = 0) terms into a proportionality constant since we are primarily interested in the shape of Σ_p_. This can be rewritten in matrix notation as

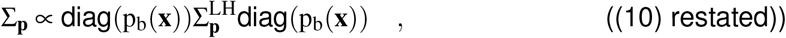

where 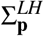 is the covariance *of the likelihood* with *s* = 0 and is task-independent. The prior, p_b_(**x**|*π* = ½)), is task-dependent. Equation (10) thus gives, to a first approximation, an expression for how noise in the likelihood is sculpted by learning: the “intrinsic” covariance in the likelihood, which is present before learning, is pre-and post-multiplied by a diagonal matrix of the task-dependent prior mass vector.

One way to reason about (10) is by considering its eigenvector decomposition. For instance, *differential correlations* are introduced only to the extent that the relative variance in the 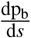 direction is increased after left-and right-multiplying the intrinsic noise 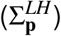 by the diagonal matrix of prior probabilities. It is nontrivial, however, to state this in terms of conditions on **x**, *s*, or 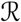, which we leave as a problem for future work.

Figure 5 was created by simulating a discretized 2D space. The likelihood functions were 2D Gaussians parameterized by *s*, so there were five degrees of freedom for each likelihood function: {*μ*_1_, *μ*_2_, *σ*_1_, *σ*_2_, *c*}, where 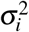 is the variance along dimension i and c is the correlation. In the first simulation, the means were parameterized by a smooth (cubic) function of *s*,

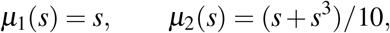

while the other three parameters did not depend on *s*. In the second simulation, means were constant while the variances and correlation were parameterized by *s* as follows:

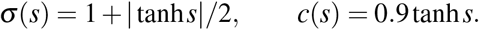

In both cases, p_e_(*s*) was set to a uniform distribution in [−3, +3]. Gaussian noise with σ = ½ was added to the means, and noise was added to the covariance of the likelihood by adding to it a random covariance matrix whose diagonal (variances) was exponential random variables and whose correlation was a tanh function of a Gaussian random variable. Starting with a uniform prior over this space, learning consisted of drawing a large number of random likelihoods (randomizing both *s* and *ε*) to estimate the average posterior, then the prior was updated to equal the average posterior, mixed with 1% of uniform density added everywhere for regularization. This process was then run to convergence in 50 independent runs of each simulation. To measure the change in covariance of the posterior density itself along dp_b_/d*s*, we compared the first and last iteration, which have the same statistics of variable likelihoods but different priors. We plotted the change in relative variance along dp_b_/d*s* in Figure 5e,j, defined as

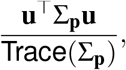

where **u** is the unit vector pointing in the dp_b_/d*s*-direction. We computed dp_b_/d*s* separately before and after learning (Figure 5d+i show dp_b_/d*s* after learning) by drawing a large number of random posteriors and taking the difference of their average at *s* = +.05 and *s* = -.05.

## Acknowledgements

We thank the many colleagues with whom we have discussed this work and who have provided us with valuable feedback, in particular (alphabetically) Matthias Bethge, Adrian Bondy, Bruce Cumming, Alex Ecker, Jakob Macke, Ruben Moreno-Bote, Hendrikje Nienborg, Maneesh Sahani, and Emmett Wyman.

This work was supported by NEI/NIH awards R01 EY028811-01 (RMH) and T32 EY007125 (RDL), as well as an NSF/NRT graduate training grant NSF-1449828 (RDL).

## Author contributions

RMH conceived the theory. RDL formalized the theory and implemented the simulations. RDL and RMH wrote the manuscript.

## Supplemental Figures

**Figure S1.**
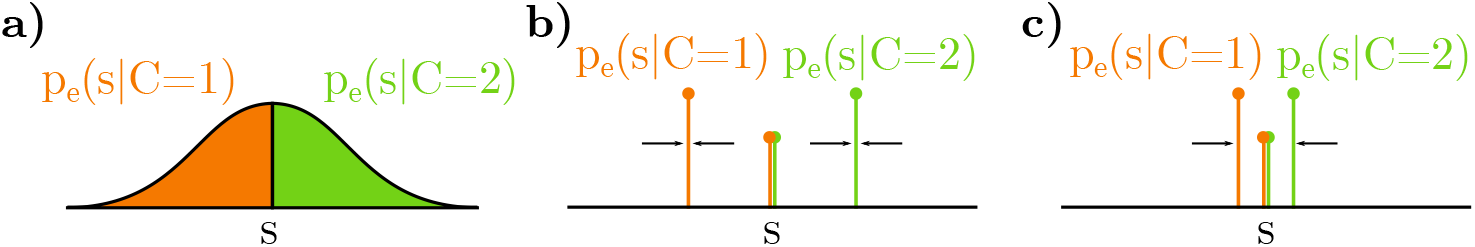
Visualizing the limiting process(es) of stimulus distributions as defined by equations (17) and (18). **a)** Initially, the distribution on stimuli may be wide, here illustrated as a Gaussian that is split by the two categories. **b)** Equation (17) considers the case where *each* category goes to a Dirac delta around some ±Δ*s*, plus a delta at zero. **c)** As the magnitude of Δ*s* gets small, the approximation in (5) gets better. As discussed in the methods, this limit may not be taken fully to Δ*s* → 0.

**Figure S2.**
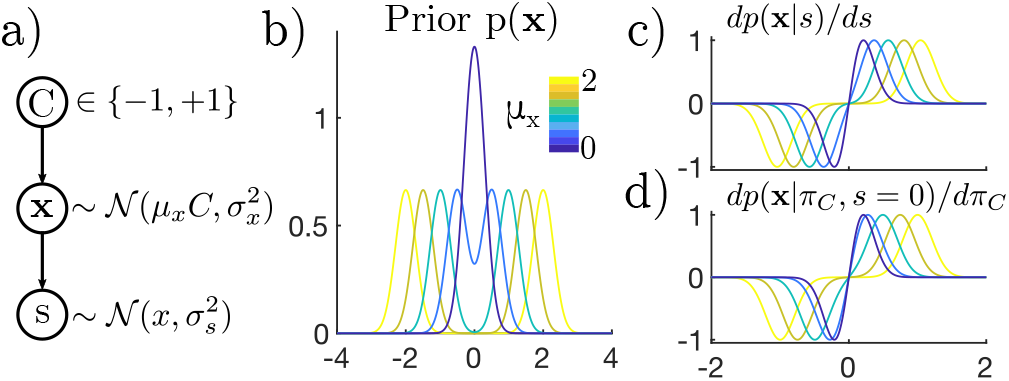
**a)** Simple generative model simulated in **b-d. x** is a scalar drawn from a Gaussian around ±*μ_x_* (matching the sign of *C*), and the stimulus *s* is drawn from a Gaussian around **x. b)** The prior on **x** is a mixture of two Gaussians. Colors correspond to different values of *μ_x_*. **c)** Derivatives of the posterior with respect to *s*. **d)** Derivatives of the posterior with respect to *π*. The match to **c** improves as *μ_x_* gets closer to 0, which simulates changes to the learned model as stimulus categories *μ_x_* draw closer together (as in Figure S1c).

**Figure S3.**
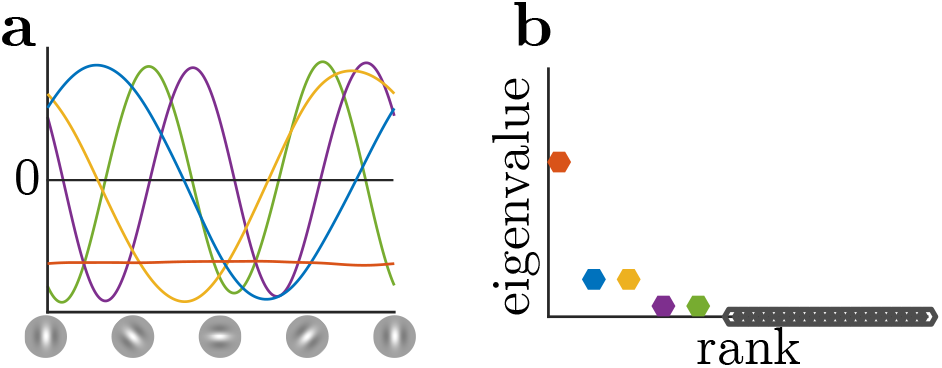
Principal components of model neurons due to only stimulus-driven correlations. Note that the sinusoidal eigenvectors at the same frequency have indistinguishable eigenvalues and hence form quadrature pairs, implying circular symmetry with respect to neurons’ tuning. There is no more variance along the vertical-horizontal preferred orientation axis than then oblique axis.

## Supplemental Text

### Note on the circularity of the ideal learning condition

Equation (2) defines the optimal task-prior (left hand side) in terms of the average posterior seen in the task (right hand side). Each posterior is, circularly, defined in terms of the prior:

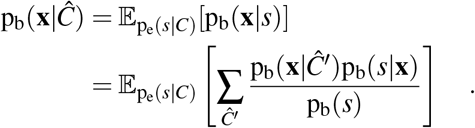

One interpretation is that equation (2) describes the *end result* of learning a task in terms of a fixed-point relation where the average posterior in the task is equal to the prior, but it does not prescribe how to arrive at such a prior.

A straightforward method to learn such a prior is to iterate until convergence, where in each step of the iteration, the “new” prior is defined as the average posterior under inferences made using the “old” prior:

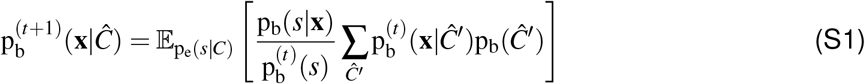

where we have assumed that it is only the prior influcence of the category on the sensory representation 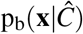, not the sensory generative procedure p_b_(*s*|**x**) that changes with learning. It follows that the the full prior on 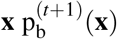 is also defined iteratively as

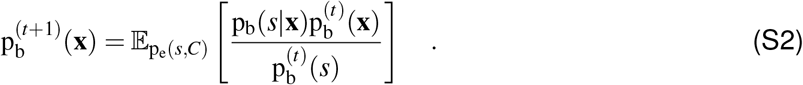

This is the iterative learning procedure used in our simulations for Figure 5.

The iterative procedure defined by equation (*S*2) has a fixed point in which the marginal likelihood on stimuli p_b_(*s*) equals the experimental distribution of stimuli p_e_(*s*), as we now show. A fixed point is reached when there is no change in the prior from one iteration to the next, so that 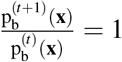.

Dividing both sides of equation (S2) by 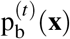 gives

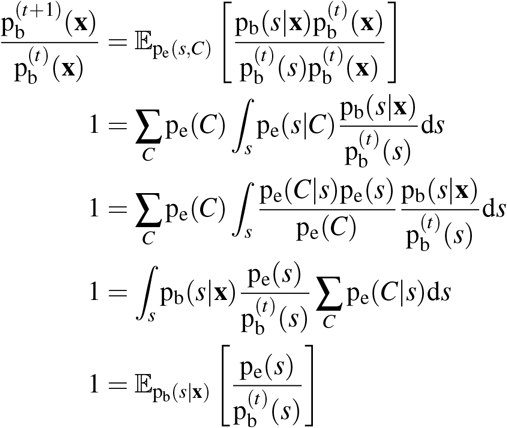

If the marginal distribution of *s* in the brain’s model at time *t* equals the experimenter’s distribution on *s*, then the term inside the expectation is 1 and hence the brain has correctly converged to a model of the task.

What we have shown here is that the apparent circularity of equation (2) is in fact a feature of any “well-calibrated” probabilistic model. The fixed-point derivation above shows that when the marginal distribution of stimuli under the brain’s (implicit) generative model matches the true distribution of stimuli defined by the experimenter, the process has converged and the relation in (2) will hold.

### Note on relaxing the limits on the stimulus distribution

Our proof of (5) required a set of two limits in which (1) the stimulus distribution approaches a mixture of Dirac deltas at *s* = 0 and *s* = ±Δ*s*, and (2) the spread ofthese components becomes small, i.e. Δ*s* gets small (but must not reach 0). These conditions might be considered extreme even for threshold psychophysics. In principle, this limits the applicability of our result whenever the empirical stimulus distribution has appreciable variance. In practice, however, three factors aid in the generality of our results. First, the stimulus distribution may be wider in the case of Linear Distributional Codes (LDCs) without affecting affecting our results for the same reason that LDCs make the same predictions in the presence of external noise. However, this would additionally require **f′** to be defined as the difference in average neural response to all stimuli in each category, by analogy to equation (23). As stated in the main text, our exact results for LDCs can be expected to degrade smoothly for nearly-linear codes.

Second, we have considered only the case where the forms a binary categorical judgment about, rather than an intermediate continuous estimation of the stimulus *s*. Even in two-alternative forced-choice tasks, subjects may internally categorize stimuli according to more than two subjective categories, for instance distinguishing “faintly rightward” separately from “strongly rightward.” To the extent that subjects *internally* make fine categorical distinctions such as this, our result for concerns categorical beliefs about “faint” categories near the *s* = 0 boundary. This necessarily involves a small range of values of *s* around *s* = 0, as in the limiting case our proof requires. Another way to say this is that forming a continuous internal *estimate* of *s* that then informs the category judgment could be formalized as a limit where the number of fine-grained categories grows large. It is, in fact, unsurprising that fluctuating internal continuous *estimates* of *s* elicit differential correlations. The limit required for our result for variable categorical beliefs can be interpreted as approaching continuous estimates around *s* = 0.

The third factor regarding generality is that the brain cannot represent arbitrary distributions, but is necessarily restricted to some finite approximation (whether by finitely many parameters in a parametric approximation, or finitely many values of **x** in a sampling-based approximation). Any family of approximations is a subspace of all possible distributions. Geometrically, one may think of “projecting” the true distributions p(**x**|…) into this subspace of approximating distributions. This projection operation will tend not to amplify differences between distributions, but will generally suppress them; the difference between approximate distributions will be less than the difference in the full space of distributions. Recall that in our derivations we used two distinct limiting processes: one where the entropy of each category shrunk (Figure S1b), and a second where their means moved towards zero (Figure S1c). After taking the first limit, the proportionality in (5) reduced to the question of whether p_b_(**x**|**E**(*s* = 0)) approximately equals 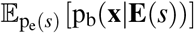. While these terms may differ significantly in probability space, their projections may not. In other words, *the brain’s distributional coding scheme may not be sensitive to these exact differences*. This suggests that the simpler the distributions represented by the brain the better our results will hold, since more distributions in the full space map to the same point in the subspace of approximate distributions when the approximating family is limited.

Taken together, these points suggest that although the proportionality in (5) is approximate, its accuracy degrades gracefully under more realistic assumptions.

### Derivation of (28) in terms of tuning to noise

If we approximate *ε* as Gaussian, then from the Taylor expansion of *f_i_*(*s* = 0, *π* = ½;*ε*) around the mean noise value, it is easy to show that the covariance between neurons *i* and *j* due to noise is approximately

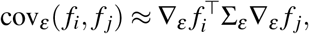

where Σ_*ε*_ is the covariance of *ε*, and ∇*_ε_f_i_* is the sensitivity of neuron *i* to variations in the noise around its mean. Computationally, the noise *ε* acts on *f_i_* through the intermediate step of the posterior, p_b_(**x**|*s* = 0, *π* = ½;*ε*). Applying the chain rule, the gradient of *f_i_* with respect to *ε* can thus be written as the product of *f_i_*’s sensitivity to p_b_(**x**|…) and the derivative of p_b_(**x**|…) with respect to *ε*. The chain rule gives 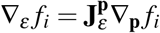, where 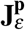 is the Jacobian (i.e. columns of **J** are gradients of elements of p_b_(**x**|…) with respect to the vector *ε*). The above covariance expression then becomes

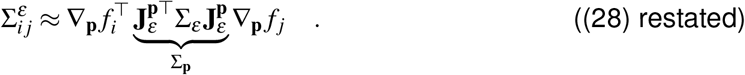

Thus we see that the covariance in neural responses induced by task-independent noise can be thought of in a two-step process: the the covariance structure of the noise (Σ_*ε*_) induces correlated variability in the posterior density (Σ_p_) through the Jacobian matrix of sensitivities 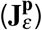, which in turn manifests as correlated *neural variability* as per the “chain rule” argument from (1).

## Footnotes

1 The term “prior” is often overloaded, referring sometimes to stationary statistics learned over long time scales, and sometimes to dynamic changes to the posterior due to higher-level inferences or internal states. Therefore, we refer to the dynamic effect of internal states on **x** as “expectations”.

2 For now we are suppressing “noise” for the sake of exposition, but will return to it later in the results.

3 Note that this excludes the possibility of separately encoding the likelihood and the prior.

## References

Aitchison L., Hennequin G., and Lengyel M. (2018). Sampling-based probabilistic inference emerges from learning in neural circuits with a cost on reliability. arXiv pp. 1–31.

Aitchson L., and Lengyel M. (2016). The Hamiltonian Brain: Efficient Probabilistic Inference with Excitatory-Inhibitory Neural Circuit Dynamics. PLoS Computational Biology pp. 1–24.

Albright T.D. (2012). On the Perception of Probable Things: Neural Substrates of Associative Memory, Imagery, and Perception. Neuron 74, 227–245.

Anderson C.H., and Van Essen D.C. (1994). Neurobiological computational systems. IEEE World Congress on Computational Intelligence pp. 1–11.

Archer E.W., Köster U., Pillow J.W., and Macke J.H. (2014). Low-dimensional models of neural population activity in sensory cortical circuits. Advances in Neural Information Processing Systems 27, 343–351.

Averbeck B.B., Latham P.E., and Pouget A. (2006). Neural correlations, population coding and computation. Nature Reviews Neuroscience 7, 358–366.

Bányai M., Lazar A., Klein L., Klon-Lipok J., Stippinger M., Singer W., and Orbán G. (2019). Stimulus complexity shapes response correlations in primary visual cortex. Proceedings of the National Academy of Sciences 116, 2723–2732.

Bányai M., and Orbán G. (2019). Noise correlations and perceptual inference. Current Opinion in Neurobiology 58, 209–217.

Beck J.M., Heller K., and Pouget A. (2013). Complex Inference in Neural Circuit swith Probabilistic Population Codes and Topic Models. Advances in Neural Information Processing Systems 25, 3068–3076.

Beck J.M., Latham P.E., and Pouget A. (2011). Marginalization in neural circuits with divisive normalization. J. Neurosci. 31, 15310–15319.

Beck J.M., Ma W.J., Kiani R., Hanks T., Churchland A.K., Roitman J., Shadlen M.N., Latham P.E., and Pouget A. (2008). Probabilistic population codes for Bayesian decision making. Neuron 60, 1142–1152.

Beck J.M., Ma W.J., Pitkow X., Latham P.E., and Pouget A. (2012). Not noisy, just wrong: the role of suboptimal inference in behavioral variability. Neuron 74, 30–39.

Berkes P., Orbán G., Lengyel M., and Fiser J. (2011). Spontaneous Cortical Activity Reveals Hallmarks of an Optimal Internal Model of the Environment. Science 331, 83–87.

Bondy A.G., Haefner R.M., and Cumming B.G. (2018). Feedback determines the structure of correlated variability in primary visual cortex. Nature Neuroscience 21, 598–606.

Bornschein J., Henniges M., and Lücke J. (2013). Are V1 Simple Cells Optimized for Visual Occlusions?AComparativeStudy. PLoSComputationalBiology 9.

Buesing L., Bill J., Nessler B., and Maass W. (2011). Neural dynamics as sampling: a model for stochastic computation in recurrent networks of spiking neurons. PLoS Computational Biology 7.

Chicharro D., Panzeri S., and Haefner R.M. (2017). Decision-related signals in the presence of nonzero signal stimuli, internal bias, and feedback. bioRxiv pp. 1–48.

Cohen M.R., and Newsome W.T. (2008). Context-Dependent Changes in Functional Circuitry in Visual Area MT. Neuron 60, 162–173.

Cunningham J.P., and Yu B.M. (2014). Dimensionality reduction for large-scale neural recordings. Nature Neuroscience 17, 1500–1509.

Dayan P., and Abbott L.F. (2001). Theoretical Neuroscience: Computational and Mathematical Modeling of Neural Systems (London: MIT Press).

de Lange F.P., Heilbron M., and Kok P. (2018). How Do Expectations Shape Perception? Trends in Cognitive Sciences 22, 764–779.

Doiron B., Litwin-kumar A., Rosenbaum R., Ocker G.K., and Josic K. (2016). The mechanics of state-dependent neural correlations. Nature Neuroscience 19, 383–393.

Echeveste R., Aitchison L., Hennequin G., and Lengyel M. (2019). Cortical-like dynamics in recurrent circuits optimized for sampling-based probabilistic inference. bioRxiv p. 696088.

Ecker A.S., Berens P., Cotton R.J., Subramaniyan M., Denfield G.H., Cadwell C.R., Smirnakis S.M., Bethge M., and Tolias A.S. (2014). State dependence of noise correlations in macaque primary visual cortex. Neuron 82, 235–248.

Ecker A.S., Berens P., Tolias A.S., and Bethge M. (2011). The Effect of Noise Correlations in Populations of Diversely Tuned Neurons. Journal of Neuroscience 31, 14272–14283.

Ecker A.S., Denfield G.H., Bethge M., and Tolias A.S. (2016). On the structure of population activity under fluctuations in attentional state. Journal of Neuroscience 0, 1–21.

Faisal A.A., Selen L.P.J., and Wolpert D.M. (2008). Noise in the nervous system. Nature Reviews Neuroscience 9, 292–303.

Felleman D.J., and Van Essen D.C. (1991). Distributed hierachical processing in the primate cerebral cortex. Cerebral Cortex 1, 1–47.

Finke R.A. (1980). Levels of equivalence in imagery and perception. Psychological Review 87, 113–132.

Fischer J., and Whitney D. (2014). Serial dependence in visual perception. Nature Neuroscience 17, 738–743.

Fiser J., Berkes P., Orbán G., and Lengyel M. (2010). Statistically optimal perception and learning: from behavior to neural representations. Trends in Cognitive Sciences 14, 119–30.

Fründ I., Wichmann F., and Macke J. (2014). Quantifying the effect of intertrial dependence on perceptual decisions. Journal of vision 14, 1–16.

Ganguli D., and Simoncelli E.P. (2014). Efficient sensory encoding and Bayesian inference with heterogeneous neural populations. Neural Computation 26, 2103–2134.

Gershman S.J., and Beck J.M. (2016). Complex Probabilistic Inference: From Cognition to Neural Computation. In Computational Models of Brain and Behavior, A. Moustafa, ed. (Wiley-Blackwell), pp. 1–17.

Gold J.I., and Shadlen M.N. (2007). The neural basis of decision making. Annual review of neuroscience 30, 535–574.

Goris R.L.T., Movshon J.A., and Simoncelli E.P. (2014). Partitioning neuronal variability. Nature Neuroscience 17, 858–865.

Green D.M., and Swets J.A. (1966). Signal Detection Theory and Psychophysics (New York: Wiley).

Haefner R.M., Berkes P., and Fiser J. (2016). Perceptual Decision-Making as Probabilistic Inference by Neural Sampling. Neuron 90, 649–660.

Haefner R.M., Gerwinn S., Macke J.H., and Bethge M. (2013). Inferring decoding strategies from choice probabilities in the presence of correlated variability. Nature Neuroscience 16, 235–242.

Haimerl C., Savin C., and Simoncelli E.P. (2019). Flexible information routing in neural populations through stochastic comodulation. Advances in Neural Information Processing Systems 33.

Hensch T.K. (2005). Critical period plasticity in local cortical circuits. Nature Reviews Neuroscience 6, 877–888.

Houlsby N.M.T., Huszár F., Ghassemi M.M., Orbán G., Wolpert D.M., and Lengyel M. (2013). Cognitive Tomography Reveals Complex, Task-Independent Mental Representations. Current Biology 23, 2169–2175.

Hoyer P.O., and Hyvärinen A. (2003). Interpreting neural response variability as monte carlo sampling of the posterior. Advances in neural information processing systems 17, 293–300.

Huang C., Ruff D.A., Pyle R., Rosenbaum R., Cohen M.R., and Doiron B. (2019). Circuit Models of Low-Dimensional Shared Variability in Cortical Networks. Neuron 101, 337–348. e4.

Kanitscheider I., Coen-Cagli R., and Pouget A. (2015). Origin of information-limiting noise correlations. Proceedings of the National Academy of Sciences 112, 6973–82.

Kersten D., Mamassian P., and Yuille A. (2004). Object perception as bayesian inference. Annual Review of Psychology pp. 271–304.

Knill D.C., and Pouget A. (2004). The Bayesian brain: the role of uncertainty in neural coding and computation. Trends in Neurosciences 27, 712–9.

Kobak D., Brendel W., Constantinidis C., Feierstein C.E., Kepecs A., Mainen Z.F., Qi X.L., Romo R., Uchida N., and Machens C.K. (2016). Demixed principal component analysis of neural population data. eLife 5, 1–36.

Kohn A., Coen-Cagli R., Kanitscheider I., and Pouget A. (2016). Correlations and Neuronal Population Information. Annual Review of Neuroscience 39, 237–256.

Körding K.P., Beierholm U.R., Ma W.J., Quartz S.R., Tenenbaum J.B., and Shams L. (2007). Causal inference in multisensory perception. PLoSOne 2.

Lange R.D., and Haefner R.M. (2017). Characterizing and interpreting the influence of internal variables on sensory activity. Current Opinion in Neurobiology 46, 84–89.

Law C.T., and Gold J.I. (2008). Neural correlates of perceptual learning in a sensory-motor, but not a sensory, cortical area. Nature Neuroscience 11, 505–513.

Law C.T.T., and Gold J.I. (2009). Reinforcement learning can account for associative and perceptual learning on a visual-decision task. Nature Neuroscience 12, 655–63.

Lee D.D., Ortega P.A., and Stocker A. (2014). Dynamic belief state representations. Current opinion in neurobiology 25, 221–7.

Lee T.S., and Mumford D. (2003). Hierarchical Bayesian inference in the visual cortex. Journal of the Optical Society of America A 20, 1434–1448.

Li N., and DiCarlo J.J. (2008). Unsupervised natural experience rapidly alters invariant object representation in visual cortex. Science 321, 1502–1507.

Lueckmann J.M., Macke J.H., and Nienborg H. (2018). Can serial dependencies in choices and neural activity explain choice probabilities? The Journal of Neuroscience 38, 2225–17.

Ma W.J., Beck J.M., Latham P.E., and Pouget A. (2006). Bayesian inference with probabilistic population codes. Nature Neuroscience 9, 1432–1438.

Ma W.J., and Jazayeri M. (2014). Neural coding of uncertainty and probability. Annual review of neuroscience 37, 205–220.

Macke J.H., and Nienborg H. (2019). Choice (-history) correlations in sensory cortex: cause or consequence? Current Opinion in Neurobiology 58, 148–154.

Marr D. (1982). Vision: A Computational Investigation into the Human Representation and Processing of Visual Information. Phenomenology and the Cognitive Sciences 8, 397.

Moreno-Bote R., Beck J.M., Kanitscheider I., Pitkow X., Latham P., and Pouget A. (2014). Information-limiting correlations. Nature Neuroscience 17, 1410–1417.

Mumford D. (1992). On the computational architecture of the neocortex. Biological cybernetics 251, 241–251.

Ni A.M., Ruff D.A., Alberts J.J., Symmonds J., and Cohen M.R. (2018). Learning and attention reveal a general relationship between neuronal variability and perception. Science 359, 463–465.

Nienborg H., Cohen M.R., and Cumming B.G. (2012). Decision-Related Activity in Sensory Neurons: Correlations Among Neurons and with Behavior. Annual Review of Neuroscience 35, 463–483.

Nienborg H., and Cumming B.G. (2007). Psychophysically measured task strategy for disparity discrimination is reflected in V2 neurons. Nature Neuroscience 10, 1608–14.

Nienborg H., and Cumming B.G. (2009). Decision-related activity in sensory neurons reflects more than a neuron’s causal effect. Nature 459, 89–92.

Nienborg H., and Cumming B.G. (2010). Correlations between the activity of sensory neurons and behavior: How much do they tell us about a neuron’s causality? Current Opinion in Neurobiology 20, 376–381.

Nienborg H., and Cumming B.G. (2014). Decision-Related Activity in Sensory Neurons May Depend on the Columnar Architecture of Cerebral Cortex. Journal of Neuroscience 34, 3579–3585.

Nienborg H., and Roelfsema P.R. (2015). Belief states as a framework to explain extra-retinal influences in visual cortex. Current opinion in neurobiology 32, 45–52.

Olshausen B.A., and Field D.J. (1996). Emergence of simple-cell receptive field properties by learning a sparse code for natural images. Nature 381, 607–609.

Olshausen B.A., and Field D.J. (1997). Sparse coding with an incomplete basis set: a strategy employed by V1?

Oram M.W., Földiák P., Perrett D.I., and Sengpiel F. (1998). The ‘Ideal Homunculus’: decoding neural population signals. Trends in Neurosciences 21, 259–265.

Orbán G., Berkes P., Fiser J., and Lengyel M. (2016). Neural Variability and Sampling-Based Probabilistic Representations in the Visual Cortex. Neuron 92, 530–543.

Parker A.J., and Newsome W.T. (1998). Sense and the single neuron: probing the physiology of perception. Annu Rev Neurosci 21, 227–277.

Pecevski D., Buesing L., and Maass W. (2011). Probabilistic inference in general graphical models through sampling in stochastic networks of spiking neurons. PLoS computational biology 7.

Petrovici M.A., Bill J., Bytschok I., Schemmel J., and Meier K. (2016). Stochastic inference with spiking neurons in the high-conductance state. Physical Review E 94.

Pitkow X., and Angelaki D.E. (2017). Inference in the Brain: Statistics Flowing in Redundant Population Codes. Neuron Perspective 94, 943–953.

Pitkow X., Liu S., Angelaki D.E., DeAngelis G.C., and Pouget A. (2015). How Can Single Sensory Neurons Predict Behavior? Neuron 87, 411–423.

Pouget A., Beck J.M., Ma W.J., and Latham P.E. (2013). Probabilistic brains: knowns and unknowns. Nature Reviews Neuroscience 16, 1170–1178.

Rabinowitz N.C., Goris R.L., Cohen M.R., and Simoncelli E.P. (2015). Attention stabilizes the shared gain of V4 populations. eLife 4.

Raju R.V., and Pitkow X. (2016). Inference by Reparameterization in Neural Population Codes. Advances in Neural Information Processing Systems 30.

Ramalingam N., McManus J.N.J., Li W., and Gilbert C.D. (2013). Top-Down Modulation of Lateral Interactions in Visual Cortex. Journal of Neuroscience 33, 1773–1789.

Rezende D.J., Mohamed S., and Wierstra D. (2014). Stochastic backpropagation and approximate inference in deep generative models. Proceedings of The 31st… 32, 1278–1286.

Ruff D.A., Ni A.M., and Cohen M.R. (2018). Cognition as a Window into Neuronal Population Space. Annual Review of Neuroscience 41, 77–97.

Sahani M., and Dayan P. (2003). Doubly Distributional Population Codes:. Neural Computation 2279, 2255–2279.

Savin C., and Denève S. (2014). Spatio-temporal representations of uncertainty in spiking neural networks. Advances in Neural Information Processing Systems 27, 1–9.

Schwartz O., and Simoncelli E.P. (2001). Natural signal statistics and sensory gain control. Nature Neuroscience 4, 819–825.

Shadlen M.N., Britten K.H., Newsome W.T., and Movshon J.A. (1996). A computational analysis of the relationship between neuronal and behavioral responses to visual motion. Journal of Neuroscience 16, 1486–1510.

Shivkumar S., Lange R.D., Chattoraj A., and Haefner R.M. (2018). A probabilistic population code based on neural samples. NeurIPS 31, 7070–7079.

Stocker A.A., and Simoncelli E.P. (2006). Noise characteristics and prior expectations in human visual speed perception. Nature Neuroscience 9, 578–585.

Stocker A.A., and Simoncelli E.P. (2007). A Bayesian Model of Conditioned Perception. Advances in NeuralInfromation Processing Systems 2007, 1409–1416.

Summerfield C., and de Lange F.P. (2014). Expectation in perceptual decision making: neural and computational mechanisms. Nature Reviews Neuroscience 15, 745–756.

Tajima C.I., Tajima S., Koida K., Komatsu H., Aihara K., and Suzuki H. (2016). Population code dynamics in categorical perception. Nature Scientific Reports 5, 1–13.

Vertes E., and Sahani M. (2018). Flexible and accurate inference and learning for deep generative models. Neural Information Processing Systems 31.

von der Heydt R., Peterhans E., and Baumgartner G. (1984). Illusory Contours and Cortical Neuron Responses. Science 224, 1260–2.

von Helmholtz H. (1925). Treatise on Physiological Optics (The Optical Society of America).

Walker E.Y., Cotton R.J., Ma W.J., and Tolias A.S. (2019). A neural basis of probabilistic computation in visual cortex. Nature Neuroscience 23, 122–129.

Wei X.X., and Stocker A.A. (2015). A Bayesian observer model constrained by efficient coding can explain ‘anti-Bayesian’ percepts. Nature Neuroscience 18, 1509–17.

Wimmer K., Compte A., Roxin A., Peixoto D., Renart A., and Rocha J.D. (2015). Sensory integration dynamics in a hierarchical network explains choice probabilities in cortical area MT. Nature Communications 6, 1–13.

Yu A.J., and Cohen J.D. (2009). Sequential effects: Superstition or rational behavior? Advances in Neural Information Processing Systems 22, 1873–80.

Yuille A., and Kersten D. (2006). Vision as Bayesian inference: analysis by synthesis? Trends in Cognitive Sciences 10, 301–8.

Zemel R.S., Dayan P., and Pouget A. (1998). Probabilistic Interpretation of Population Codes. Neural Computation 10, 403–430.

Zohary E., Shadlen M.N., and Newsome W.T. (1994). Correlated Neuronal Discharge rate and its implications for psychophysical performance. Letters to Nature 370, 140–143.

